# Gaze-stabilizing central vestibular neurons project asymmetrically to extraocular motoneuron pools

**DOI:** 10.1101/151548

**Authors:** David Schoppik, Isaac H. Bianco, David A. Prober, Adam D. Douglass, Drew N. Robson, Jennifer M.B. Li, Joel S.F. Greenwood, Edward Soucy, Florian Engert, Alexander F. Schier

## Abstract

Within reflex circuits, specific anatomical projections allow central neurons to relay sensations to effectors that generate movements. A major challenge is to relate anatomical features of central neural populations — such as asymmetric connectivity — to the computations the populations perform. To address this problem, we mapped the anatomy, modeled the function, and discovered a new behavioral role for a genetically-defined population of central vestibular neurons in rhombomeres 5-7 of larval zebrafish. First, we found that neurons within this central population project preferentially to motoneurons that move the eyes downward. Concor-dantly, when the entire population of asymmetrically-projecting neurons was stimulated collectively, only downward eye rotations were observed, demonstrating a functional correlate of the anatomical bias. When these neurons are ablated, fish failed to rotate their eyes following either nose-up or nose-down body tilts. This asymmetrically-projecting central population thus participates in both up and downward gaze stabilization. In addition to projecting to motoneurons, central vestibular neurons also receive direct sensory input from peripheral afferents. To infer whether asymmetric projections can facilitate sensory encoding or motor output, we modeled differentially-projecting sets of central vestibular neurons. Whereas motor command strength was independent of projection allocation, asymmetric projections enabled more accurate representation of nose-up stimuli. The model shows how asymmetric connectivity could enhance the representation of imbalance during nose-up postures while preserving gaze-stabilization performance. Finally, we found that central vestibular neurons were necessary for a vital behavior requiring maintenance of a nose-up posture: swim bladder inflation. These observations suggest that asymmetric connectivity in the vestibular system facilitates representation of ethologically-relevant stimuli without compromising reflexive behavior.

**Significance Statement:** Interneuron populations use specific anatomical projections to transform sensations into reflexive actions. Here we examined how the anatomical composition of a genetically-defined population of balance interneurons in the larval zebrafish relates to the computations it performs. First, we found that the population of interneurons that stabilize gaze preferentially project to motoneurons that move the eyes downward. Next, we discovered through modeling that such projection patterns can enhance the encoding of nose-up sensations without compromising gaze stabilization. Finally we found that loss of these interneurons impairs a vital behavior, swim bladder inflation, that relies on maintaining a nose-up posture. These observations suggest that anatomical specialization permits neural circuits to represent relevant features of the environment without compromising behavior.

## Introduction

Neural circuits utilize populations of interneurons to relay sensation to downstream effectors that in turn generate behavior. The anatomical composition of interneuron populations has provided insight into its function. For example, interneuron populations are often organized into maps composed of non-uniformly sized sets of neurons similarly sensitive to particular features (Kaas, 1997). Such visual topography in the thalamus (Connolly and Essen, 1984) and cortex (Daniel and Whitteridge, 1961) magnifies the input from the central visual field. This magnification is thought to underlie enhanced perceptual acuity (Duncan and Boynton, 2003). Preferential anatomical organization is thought to facilitate adaptive olfactory (Hansson and Stensmyr, 2011), visual (Barlow, 1981; Xu et al., 2006), somatosensory (Adrian, 1941; Catania and Remple, 2002), and auditory (Bendor and Wang, 2006; Knudsen et al., 1987) computations. However, little is known about how these anatomical asymmetries within populations of sensory interneurons determine the activity of their target motor effectors. Motor anatomy shares a similar uneven organization (Kuypers, 2011), but the complex spatiotemporal encoding of muscle synergies have made comparable dissection of motor circuits more challenging (Harrison and Murphy, 2014; Levine et al., 2012; Shenoy et al., 2013). Even where descending cortical (Lemon, 2008) or brainstem (Esposito et al., 2014) neurons synapse directly on motoneurons, the complexity of most behaviors make it difficult to relate anatomy to function. Data relating the anatomical projections of interneuron populations to their function is needed to address this problem.

By virtue of their defined connectivity, interneurons within central reflex circuits offer the opportunity to explore the relationship between population-level anatomical properties and function in a simpler framework. Vestibular interneurons, an ancient and highly conserved population, transform body/head destabilization into commands for compensatory behaviors such as posture and gaze stabilization (Straka and Baker, 2013; Straka et al., 2014; Szentágothai, 1964). Gaze-stabilizing vestibular brainstem neurons receive innervation from peripheral balance afferents (Uchino et al., 2001) and use highly stereotyped axonal projections to particular extraocular motoneuron targets that produce directionally-specific eye movements (Iwamoto et al., 1990b; McCrea et al., 1987; Uchino et al., 1982). One anatomical and physiological characterization of up/down-sensitive vestibular neurons in the cat suggested a potential 3:1 bias towards neurons responsible for downward eye movements (Iwamoto et al., 1990a) However, extracellular recording experiments may be subject to selection bias. Further, as selective activation has been impossible, whether there are functional correlates of putative anatomical specialization remains unknown.

To study the relationship between the anatomical specializations of interneuron populations and their functions, we investigated a genetically-defined population of vestibular brainstem neurons in a model vertebrate, the larval zebrafish. Larval zebrafish face well-defined challenges that necessitate control of body orientation in the vertical/pitch axis (i.e. nose-up/nose-down). First, larval zebrafish rely on vestibular sensation to guide upward swimming to the water’s surface to gulp air and inflate their swim bladders, a vital organ necessary to maintain buoyancy (Goolish and Okutake, 1999; Riley and Moorman, 2000). Further, fish actively maintain a nose-up posture (Ehrlich and Schoppik, 2017), permitting them to efficiently maintain their position in the water column despite being slightly denser than their surroundings (Aleyev, 1977; Stewart and McHenry, 2010). Larval zebrafish utilize vestibular brainstem neurons to stabilize gaze by performing torsional and vertical eye movements (Bianco et al., 2012). These same neurons project to nuclei responsible for movement initiation and pitch tilts (Pavlova and Deliagina, 2002; Severi et al., 2014; Thiele et al., 2014; Wang and McLean, 2014).

We leveraged known properties of the gaze-stabilization circuit to relate the anatomy of a genetically-defined population of vestibular brainstem neurons and their function. Our study reports three major findings. First, we discovered that central vestibular neurons in rhombomeres 5-7 (R5-R7) project preferentially to extraocular motoneurons that move the eyes down. Ablation of these neurons eliminates counter-rotation of the eyes following body tilts, establishing a role in gaze-stabilization. Second, modeling revealed that asymmetrically projecting neurons could enhance the capacity to represent nose-up stimuli without compromising gaze-stabilization. Third, we discovered that fish do not inflate their swim bladders following ablation of these interneurons. Taken together, our data suggest that the anatomical specialization we observe permits sensory specialization while maintaining reflexive capabilities.

## Methods

### Fish Care

All protocols and procedures involving zebrafish were approved by the Harvard University Faculty of Arts & Sciences Standing Committee on the Use of Animals in Research and Teaching (IACUC). All larvae were raised at 28.5° C, on a standard 14/10 hour light/dark cycle at a density of no more than 20-50 fish in 25-40mL of buffered E3 (1mM HEPES added). When possible, experiments were done on the mitfa-/- background to remove pigment; alternatively, 0.003% phenylthiourea was added to the medium from 24hpf onwards and changed daily. Larvae were used from 2 days post-fertilization (dpf) to 11 dpf. During this time, zebrafish larvae have not determined their sex.

### Behavior

Torsional eye movements were measured following step tilts delivered using an apparatus similar in design to (Bianco et al., 2012). All experiments took place in the dark. Larval fish were immobilized completely in 2% low-melting temperature agar (Thermo Fisher 16520), and the left eye was freed. The agar was then pinned (0.1mm stainless minutien pins, FST) to a 5mm^2^ piece of Sylgard 184 (Dow Corning) which was itself pinned to Sylgard 184 at the bottom of a 10mm2 optical glass cuvette (Azzota, via Amazon). The cuvette was filled with 1mL of E3 and placed in a custom holder on a 5-axis (X,Y,Z,pitch,roll) manipulator (ThorLabs MT3 and GN2). The fish was aligned with the optical axes of two orthogonally placed cameras such that both the left utricle and two eyes were level with the horizon (front camera).

The eye-monitoring camera (Guppy Pro 2 F-031, Allied Vision Technologies) used a 5x objective (Olympus MPLN, 0.1 NA) and custom image-forming optics to create a 100x100 pixel image of the left eye of the fish (6 *μm*/pixel), acquired at 200Hz. The image was processed on-line by custom pattern matching software to derive an estimate of torsional angle (LabView, National Instruments), and data were analyzed using custom MATLAB scripts (Mathworks, Natick MA). A stepper motor (Oriental Motors AR98MA-N5-3) was used to rotate the platform holding the cameras and fish. The platform velocity and acceleration was measured using integrated circuits (IDG500, Invensense and ADXL335, Analog Devices) mounted together on a breakout board (Sparkfun SEN-09268). Fish were rotated stepwise for 10 cycles: from 0° to −60°, where positive values are nose-down, then from −60° to 60°, and then back to 0° in 10° increments, with a peak velocity of 35°/sec. The inter-step interval was 5 seconds, and the direction of rotation was then reversed for the next sequence of steps.

The eye’s response across the experiment was first centered to remove any offset introduced by the pattern-matching algorithm. Data were then interpolated with a cubic spline interpolation to correct for occasional transient slowdowns (i.e. missed frames) introduced by the pattern-matching algorithm. The eye’s velocity was estimated by differentiating the position trace; high-frequency noise was minimized using a 4-pole low-pass Butterworth filter (cutoff = 3Hz). Each step response was evaluated manually; trials with rapid deviations in eye position indicative of horizontal saccades or gross failure of the pattern-matching algorithm were excluded from analysis. The response to each step for a given fish was defined as the mean across all responses to that step across cycles. The gain was estimated by measuring the peak eye velocity occurring over the period 375-1000 ms after the start of the step. The steady-state response was estimated by measuring the mean eye position over the final 2 sec of the step; the range was the difference between the most eccentric nose-up and nose-down steady-state angles.

Gain was evaluated over the range from +30° to −30°, i.e. the first three steps away from the horizontal meridian. We chose this interval for three reasons: 1) Fish spend the overwhelming majority of their time with a body orientation in this range (Ehrlich and Schoppik, 2017) 2) The responses here were the strongest, allowing us confidence in the dynamic capacity of the system without encountering the biophysical limits imposed by orbital structure 3) Because the utricle conveys information both about static and dynamic changes in orientation, the eyes adopt an increasingly eccentric rotation as the stimulus progresses, potentially constraining dynamic range.

### Transgenic Lines

Tg(-6.7FRhcrtR:gal4VP16):-6.7FRhcrtR was amplified the using a nested PCR strategy. First, a 6775bp DNA fragment immediately upstream of the Fugu rubripes hcrtr2 start site was amplified from genomic DNA, using a high-fidelity polymerase (PfuUltra II Fusion, Stratagene) with primers 5’-AATCCAAATTCCCAGTGACG-3’ and 5’-CCAGATACTCGGCAAACAAA-3’, 56° C annealing temperature, 1:45 elongation time. The PCR product was TOPO cloned into a TA vector (Thermo Fisher). Using the resulting plasmid as a template, a 6732bp fragment was amplified using primers 5’-AATCCAAATTCCCAGTGACG-3’ and 5’-CCAGATACTCGGCAAACAAA-3’ 55° C annealing temperature, 1:45 elongation and similarly TOPO cloned into a GATEWAY-compatible vector (PCR8/GW, Thermo Fisher). The resulting entry vector was recombined into a destination vector upstream of gal4-VP16, between Tol2 integration arms (Urasaki et al., 2006). Tg(UAS-E1b:Kaede)s1999t embryos were injected at the one-cell stage with 0.5nL of 50ng/uL plasmid and 35ng/uL Tol2 transposase mRNA in water, and their progeny screened for fluorescence. One founder produced three fluorescent progeny; one survived. To identify transgenic fish without using a UAS reporter, potential carriers were genotyped using the following primers to generate a 592bp product spanning the upstream Tol2 arm and the start of the Fugu sequence: 5’-CAATCCTGCAGTGCTGAAAA-3’ and 5’-TGATTCATCGTGGCACAAAT-3’ 57° C annealing temperature, 0:30 elongation time. The complete expression pattern has been described elsewhere (Lacoste et al., 2015) and is part of the Z-brain atlas (Randlett et al., 2015)

Tg(14xUAS-E1b:hChR2(H134R)-EYFP):hChR2(H134R)-EYFP (Zhang et al., 2007) was subcloned down-stream of 14 copies of a UAS element and an E1b minimal promoter in a vector containing an SV40 polyA sequence and Tol2 recognition arms (Urasaki et al., 2006). This vector was co-injected with tol2 transposase mRNA into TLAB embryos at the single cell stage. Potential founders were screened by crossing to Tg(isl1:Gal4-VP16,14xUAS:Kaede)(Pan et al., 2011) and monitoring tail movements in response to blue light from an arc lamp on a stereomicroscope (Leica MZ16) at 30hpf.

The following transgenic lines were used: Tg(UAS-E1b:Kaede)s1999t (Scott et al., 2007), Tg(isl1:GFP) (Higashijima et al., 2000), Tg(UAS:KillerRed) (Bene et al., 2010), Tg(UAS-E1b:Eco.NfsB-mCherry) (Pisharath et al., 2007), atoh7th241/th241 (Kay et al., 2001); Tg(atoh7:gap43-RFP) (Zolessi et al., 2006), Tg(5xUAS:sypb-GCaMP3) (Nikolaou et al., 2012) and Et(E1b:Gal4-VP16)s1101t (Scott et al., 2007).

### Anatomy

To generate mosaically-labeled fish, 0.5nL of 30ng/*μL* plasmid DNA (14xUAS-E1b:hChR2(H134R)-EYFP (Douglass et al., 2008) or UAS-Zebrabow (Pan et al., 2013)) was injected in water at the one-cell stage into Tg(-6.7FRhcrtR:gal4VP16); Tg(isl1:GFP) fish. Embryos were screened at 24-48hpf. The majority (80%) of injected fish were excluded due to deformities or developmental arrest. The remaining fish were screened at 72hpf under a fluorescent stereoscope (Leica MZ16) with a high-pass GFP emission filter for YFP fluorescence or a Cy3 emission filter for dTomato. As Tg(-6.7FRhcrtR:gal4VP16) will label the skin and notochord early (36-48hpf) and fluorescence in either structure is relatively easy to visualize, embryos with mosaic labeling (usually 1-10 cells) in these structures were selected. On average, 1-2% of injected embryos were retained for high-resolution screening. Larvae were anesthetized (0.016% w/v tricaine methane sulfonate, Sigma A5040) mounted dorsally at 5-7dpf and imaged on a confocal microscope (Zeiss 510, 710, or 780, using either a 20x 1.0N.A., a 40x 1.1 N.A. or a 63x 1.0 N.A. objective with Zen 2010, 8-bit acquisition) with excitation of 488nm (GFP) and 514nm (EYFP), and emission for the two channels was either separated at 550nm by a glass dichroic filters or a tunable filter. The two channels could reliably be separated provided the level of EYFP was strong relative to GFP.

Most of the fish selected for confocal imaging had some neurons labeled in the brain, but on average, only 0.5%-2% (i.e. 5-20 for every 1000) of injected embryos would have vestibular nucleus neurons that were both bright and sufficiently isolated enough to trace. Neurons were only included in the study if their axon could be traced unambiguously throughout its entirety to a distinct cell body; qualitatively, the asymmetry persisted among excluded fish. Neurons were traced manually with the assistance of the ImageJ plugin Simple Neurite Tracer (Longair et al., 2011). Cell bodies of the oculomotor and trochlear nuclei were localized manually using the Fiji/ImageJ ROI functionality (Schindelin et al., 2012). Superior oblique motoneurons were found in nIV and superior rectus motoneurons were the most ventral somata in nIII (Greaney et al., 2016). All images were adjusted linearly, using the Brightness & Contrast functionality in Fiji/ImageJ (Schindelin et al., 2012). For display purposes, a non-linear histogram adjustment (gamma = 0.5) was applied to the maximum intensity projection in Figure 1b and 2a to increase the relative brightness of thin axonal arbors, and, for Figure 2a, to make clear the sparse nature of the label.

**Figure 1:**
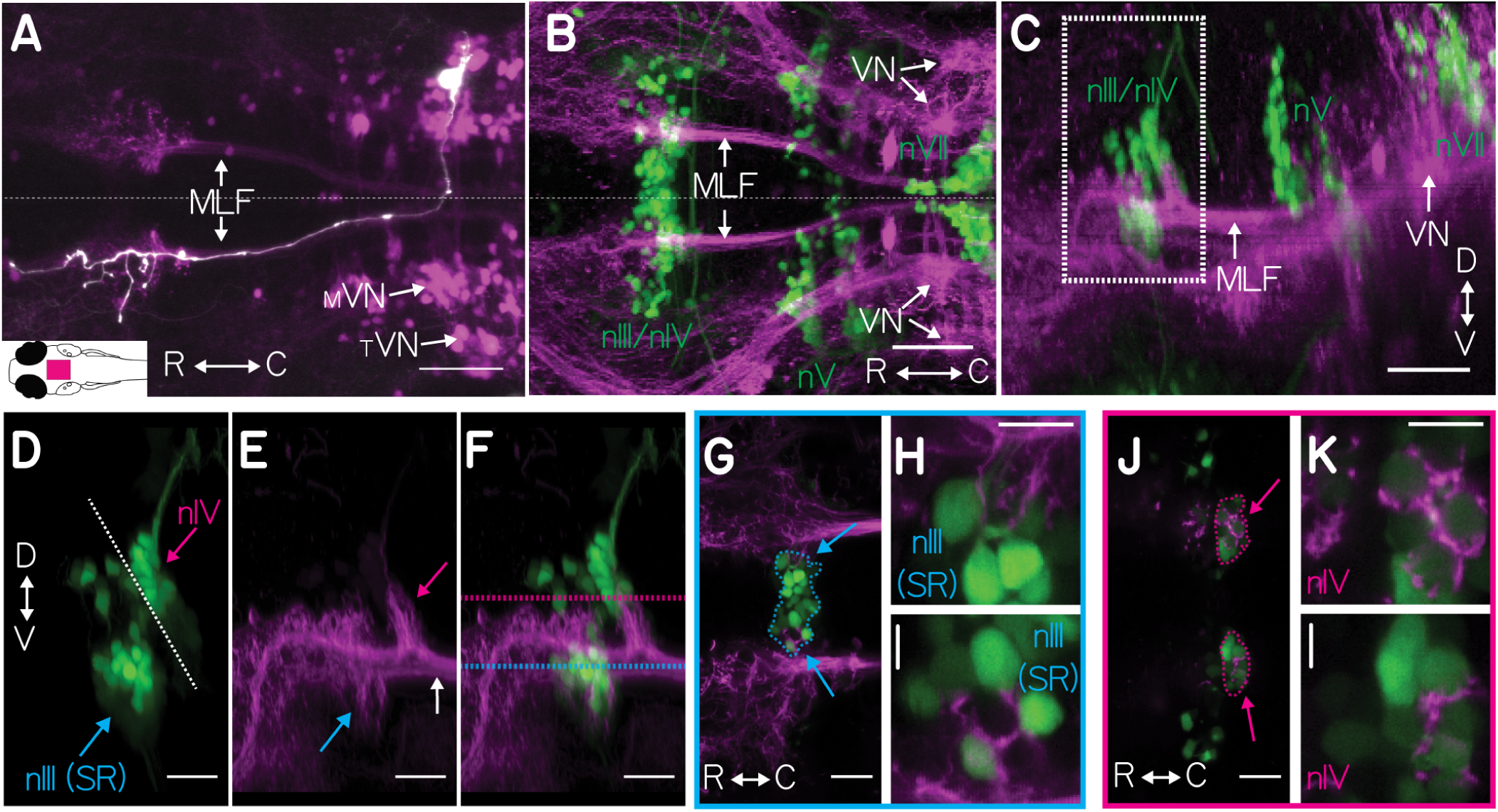
Vestibular nucleus neurons labeled in Tg(-6.7FRhcrtR:gal4VP16). A, Morphology of a single filled vestibular neuron (white) and all vestibular neurons (purple). Arrows point to the cells that belong to two sub-populations of the vestibular nuclei: the tangential (_*T*_ VN) and medial vestibular (_*M*_ VN) and the path their axons take along the medial longitudinal fasciculus (MLF). The expression pattern of Tg(-6.7FRhcrtR:gal4VP16); Tg(UAS-E1b:Kaede)s1999t (purple) is shown as a transverse maximum intensity projection (MIP), with one vestibular neuron, co-labeled by focal electroporation of gap43-EGFP (white). Inset schematic of a dorsal view of a larval zebrafish, with a magenta rectangle indicating the location of the image. Scale 50 *μm* B-C, Labeled vestibular nucleus neurons (purple) project to ocular motoneurons (green). Transverse (B) and sagittal (C) MIP of Tg(-6.7FRhcrtR:gal4VP16);Tg(UAS-KillerRed) (purple);Tg(isl1:GFP) (green, image gamma = 0.5) showing cranial motoneurons from nIII/nIV, nV, and nVII (green text). Arrows highlight the cell bodies and axons of labeled neurons in the two populations of vestibular neurons projecting along the MLF. Scale 50 *μm*. D-F, Close-up of ocular motoneuron region (white boxed region in 1C), showing major branch patterns of vestibular neuron axon fascicle (purple) relative to motoneurons (green). 1D shows motoneurons from Tg(isl1:GFP) (green) in nIV (magenta arrow), superior rectus motoneurons of nIII (cyan arrow) and the midbrain/hindbrain boundary (white dotted line). 1E shows branches of the vestibular neuron axon fascicle (purple), as they emerge from the MLF (white arrow) in Tg(- 6.7FRhcrtR:gal4VP16);Tg(UAS-KillerRed). First projection is to nIV (magenta arrow), second projection to nIII (cyan arrow). F, merge of 1D-1E. Scale 20 *μm*. G-I, Broad and close-up views of vestibular neuron axonal projection (purple) to nIII cell bodies (green), taken at the transverse plane delineated by the cyan dotted line in 1F, superior rectus (SR) motoneurons (nIII) encircled in cyan. Cyan arrows in 1G localize close-ups in 1H and 1I. Scale 10 *μm* J-L, Broad and close-up view of vestibular neuron axonal projection (purple) to nIV cell bodies (green), taken at the transverse plane delineated by the magenta dotted line in 1F, superior oblique (SO) motoneurons (nIV, green) encircled in magenta. Arrows in 1J point to close-up in 1K and 1L. Scale 10 *μm*.

**Figure 2:**
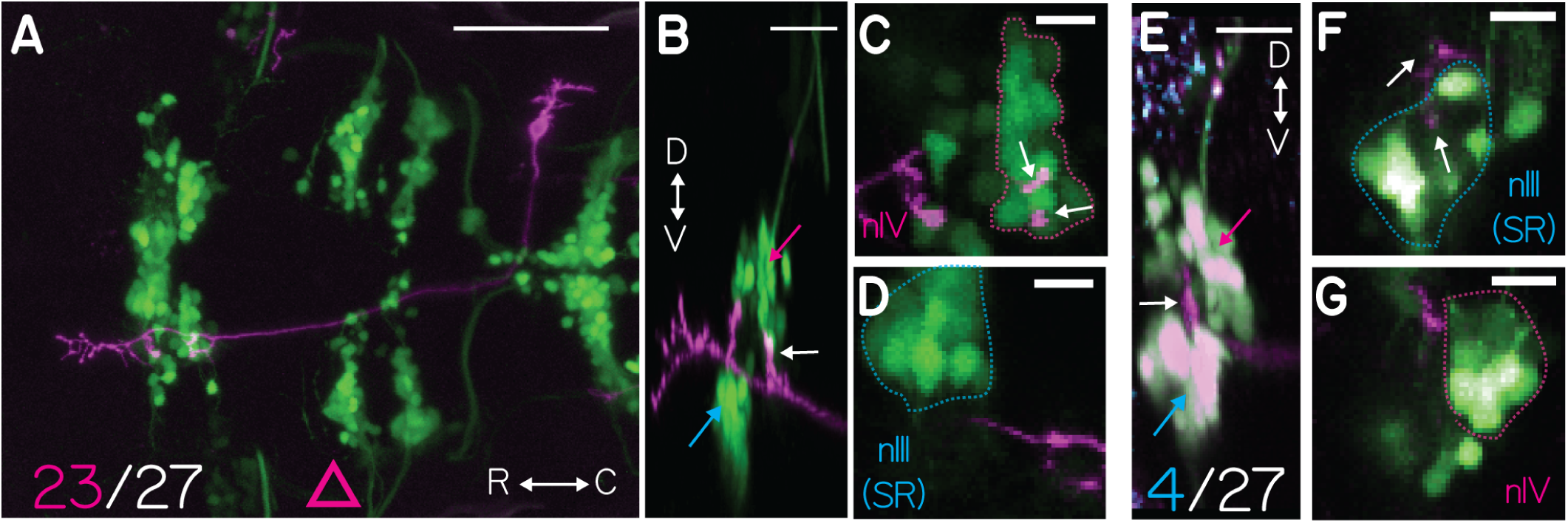
Projections from singly labeled vestibular nucleus neurons. A, Example vestibular neuron (purple) that projects to nIV (nose-up) motoneurons (green), 23/27 neurons projected to nIV. Transverse MIP of a single vestibular neuron labeled with UAS-ChR2(H134R)-EYFP (purple) in Tg(-6.7FRhcrtR:gal4VP16);Tg(isl:GFP) (green). Gamma = 0.5 to highlight the sparse label. Scale 100 *μm*. Pink triangle refers to the data in Figure 7D. B, Sagittal MIP of the neuron in Figure 2A highlighting nIII (cyan arrow), nIV (magenta arrow), and projection to nIV (white arrow). Scale 20 *μm*. C, Transverse MIP of nIV (green cell bodies in dotted magenta outline) from 2A, Vestibular neuron projection (purple axon, white arrow) Scale 10 *μm*. D, Transverse MIP of nIII (green cell bodies in dotted cyan outline) with no proximal vestibular neuron (purple) projection. E, Example vestibular neuron (purple) that projects to nIII SR (nose-down) motoneurons (green), 4/27 neurons projected exclusively to nIII. Sagittal MIP of a single axon expressing 14xUAS-E1b:hChR2(H134R)-EYFP (purple) in Tg(-6.7FRhcrtR:gal4VP16);Tg(isl1:GFP) (green); Tg(atoh7:gap43-RFP)(cyan) fish. Expression of bright EGFP bleeds into the purple channel, making the cell bodies white. nIV (magenta arrow) nIII (cyan arrow) and the vestibular neuron projection to SR motoneurons in nIII (white arrow). Scale 20 *μm*. F, Transverse MIP of nIII (cells in blue outline) from 2e, purple projections from vestibular neuron (white arrow). Scale 10 *μm* G, Transverse MIP of nIV (cells in magenta outline) from 2e with no purple vestibular neuron projection. Scale 10 *μm*.

Retrograde labeling of the ocular motor nuclei was done as previously described (Greaney et al., 2016; Ma et al., 2010). In brief, crystals of fluorescently-conjugated dextrans (10,000 MW, Thermo Fisher D-1824 or D-22914) were placed in the left orbit of anesthetized 5-7dpf fish. In fish, the superior eye muscles receive projections from the contralateral motor nuclei, making the relevant neurons in nIV (superior oblique) and nIII (superior rectus) easy to discriminate, as they were exclusively labeled on the contralateral (right) side.

Focal electroporations were done as detailed previously (Bianco et al., 2012; Tawk et al., 2009). Briefly, anesthetized larvae (2 dpf) were immobilized in low-melting temperature agarose. Micropipettes (tip diameter of 1-2 mm) were filled with a solution containing 1 mg/ml gap43-EGFP plasmid DNA in distilled water. To target the vestibular nucleus neurons, the pipette was placed at the lateral limit of rhombomere 5, using the decussation of the Mauthner axon midline crossing as a landmark. A Grass SD9 stimulator (Grass Technologies) was used to deliver three trains of voltage pulses in succession, with 1 s interval between trains. Each train was delivered at 200Hz for 250ms, 2ms on time, with an amplitude of 30V. Larvae were imaged at 5dpf on a custom multi-photon microscope at 790 nm.

### Lesions

Single-cell ablations were performed using a pulsed infrared laser (SpectraPhysics MaiTai HP) at 820nm (80MHz repetition rate, 80 fs pulse duration) at full power: 200mW (2.5nJ) measured at the specimen with a power meter (ThorLabs S130C). Fish were mounted dorsally in 2% low-melt agarose in E3 under a 20x 0.95 NA objective (Olympus) and anesthetized as described above. Cell bodies were targeted for ablation based on anatomical location, starting with the most ventro-lateral neurons in the tangential nucleus and then moving dorso-medially through the tangential and medial vestibular nucleus. Each cell was exposed to the pulsed infrared laser light for a brief period of time (35-50ms) while the resulting fluorescent emissions were measured; usually, there was a brief pulse of light that saturated the detection optics which was used to shutter the laser. 5-10 neurons/plane were targeted bilaterally, resulting in either loss of fluorescence (Tg(UAS-E1b:Kaede)s1999t and Tg(isl1:GFP)) or increased diffuse fluorescence at the cell body (Tg(UAS-ChR2-E134R-EYFP)). Fish were imaged immediately and 24 hours after ablation to confirm the extent of the lesion. 15% of lesioned fish were excluded because they did not survive a full 24hrs after the lesion. Fish were observed under a stereomicroscope in a petri dish post-lesion to ensure the presence of spontaneous horizontal saccades and normal jaw movements; all lesioned fish showed both. Fish for lesions were 4-5 dpf, as preliminary experiments showed that plasma formation was more effective in younger fish, and were selected to be the brightest in the clutch (likely doubly homozygous for UAS-E1b:Kaede and 6.7FRhcrtR:gal4VP16).

As previously described (Bianco et al., 2012), the eye movements in younger fish are of lower gain, and 3/17 fish were excluded from analysis because their total range was < 10°. Behavior was always measured at least 4 hours and no more than 8 hours after lesions. The decrease in gain was reported as a percentage of pre-lesion gain, defined as the difference between the median pre-lesion gain and median post-lesion gain normalized by the median pre-lesion gain. To activate KillerRed, green light (Zeiss set 43, 545nm/25) from an arc lamp was focused through a 63x 1.0 NA objective stopped down to fill a 200 *μm* diameter region for 15 minutes. Fish were mounted dorsally and anesthetized as described above. The focal plane was at the level of the decussation of the Mauthner axons, measured under brightfield illumination. Due to equipment replacement the precise power of the arc lamp could not be measured, but 20 minutes of exposure under identical conditions was fatal to the fish. Post-lesion behavior was measured at least 4 hours after the light exposure. To induce apoptosis with nitroreductase, fish were placed in E3 with 7.5mM of metrodinazole (Sigma M1547) in 0.2% v/v DMSO and behavior was measured 24hrs later (Curado et al., 2007). The presence of mCherry fluorescence was assayed after behavior to determine genotype.

### Optical Activation and Analysis

Channelrhodopsin-induced eye movements were monitored using the same apparatus used for measuring tilt-induced behavior, with the addition of a fiber-coupled laser on an independent micromanipulator (Arrenberg et al., 2009; Schoonheim et al., 2010). Fish were immobilized and mounted as before, and agar was removed above the head as well as the left eye. Stimulus was generated by a 100mW 473nm diode laser (Shanghai DreamLasers SDL-473-100MFL) coupled by the manufacturer to a 50 *μm* inner diameter 0.22 NA multimode fiber (ThorLabs AFS50/125Y) that itself was butt-coupled to a 10mm cannula made from the same diameter fiber (ThorLabs AFS50/125YCANNULA). Power at the cannula tip was 30-60mW, measured with a power meter (ThorLabs S130C). The fiber tip was placed above the ear, evenly-centered between the eyes, and 1mm above the skin of the fish. Stimuli ranged in duration from 1 *μsec* to 100 msec, and were presented every 5 seconds. Eye movements were tracked and processed as before, including manual analysis; only fish with at least 25 analyzable responses to a given stimulus were included in the analysis. The response to a given stimulus was quantified by taking the peak angular rotation reached over the first 2 sec.

By microinjecting plasmid DNA at the single-cell stage, we generated embryos as above with somatic expression of ChR2-EYFP in random subsets of vestibular neurons, on a blind background, atoh7th241/atoh7th241 (Kay et al., 2001). As with anatomical experiments, between 5-20 fish for each 1000 injected had acceptable expression. Of these, only 1/4 were homozygous for atoh7th241, and only 1/4 of those expressed the allele necessary to confirm blindness by visualizing the absence of retinal ganglion cell axons Tg(atoh7:gap43-RFP). The large number of alleles required and the low success rate limited the number of fish available to test. Tracing individual axonal projections to quantify the absolute number of VNs labeled in a given fish was not possible except in the most sparsely labeled fish. Further, as expected with somatic expression, ChR2-EYFP levels varied considerably across vestibular neurons. To measure the relationship between expression levels/number of labeled neurons and the magnitude of the evoked eye movement, we quantified EYFP fluorescence. Vestibular neurons are the only neurons with rostral MLF projections labeled in Tg(-6.7FRhcrtR:gal4VP16). As such, the total intensity of the MLF projection for a given fish was measured from the rostral-most point behind nIV, stopping caudally where the projection narrows to the midline (rhombomere 4). A single image that summed the intensity of all slices in the confocal stack that contained the MLF projection was used for our measurements. To correct for differences in acquisition parameters, MLF fluorescence was normalized by a measure of acquisition noise. Noise was estimated by measuring the summed fluorescence of a region between the branches of the MLF, which did not contain any neuropil. A value of one indicates no MLF fluorescence differentiable from background noise, two indicates MLF fluorescence twice that of the background, etc.. Ocular responses to blue light were evaluated and reported as above. Responses were evaluated for significance by comparing the median activity 200 ms after the stimulus to the baseline (200 ms before the stimulus).

### Model

Our model estimated the collective activity of 80 post-synaptic neurons generated by integrating activity from a set of pre-synaptic neurons. We evaluated two free parameters: the number of pre-synaptic neurons in the set (30, 42, 70, 105, 140, 168, 180) and the number of inputs on to a given post-synaptic neuron (2-30). Presynaptic activity was generated by translating a rate function, derived from the velocity profile of the steps used in the behavioral experiment, into a Poisson train of activity. Step velocity was scaled to match the reported velocity sensitivity (2 spikes/°/sec) of second-order vestibular neurons (Iwamoto et al., 1990a) to generate a rate function for Poisson spikes. The velocity reached a peak of 35 °/sec and lasted 1sec; the model was run at 1kHz. Poisson trains were subjected to an imposed 2msec refractory period. The spikes were then convolved with a decaying exponential with *τ* = 1.5sec to represent an excitatory post-synaptic potential. A random subset of pre-synaptic neurons were selected from the set and summed together to create an input to a post-synaptic neuron. Post-synaptic activity was determined by thresholding the input, subject to a 2msec refractory period. The threshold for the post-synaptic neuron was defined as the minimum of an input of 1.8 or 95% of the cumulative distribution of pre-synaptic input strength. One input spike had a value of 1; after convolution, a threshold of 1.8 was reached if at least four spikes were present across all inputs over a 4ms period. Changing the threshold ensured that the post-synaptic response would not saturate as the number of inputs increased; the specific threshold did not change the relationships we observed and is expected from the basic properties of extraocular motoneurons (Torres-Torrelo et al., 2012). We generated 80 distinct spike trains, reflecting the number of motoneurons in a given motoneuron pool (Greaney et al., 2016). The total post-synaptic response was defined as the average activity, evaluated where the rate function was positive. The strength of the relationship between the pre-synaptic rate function and the summed post-synaptic response was defined as the coefficient of determination.

### Statistics

As data were not normally distributed, expected values are reported as the median, variability as the median absolute deviation (MAD), and non-parametric tests of significance were used. Potential differences between groups (e.g. up tilts vs. down) were evaluated using the Wilcoxon rank sum test, and the Wilcoxon signed rank test was used to test whether a distribution had a median different from zero (e.g. change in performance postlesion). Significance was determined at p < 0.05.

## Results

A genetically-defined population of brainstem neurons projects preferentially to extraocular motoneurons that move the eyes downward

We adopted a molecular approach to characterize a subset of vestibular brainstem neurons in the larval zebrafish. We used a transgenic line of zebrafish, Tg(-6.7FRhcrtR:gal4VP16) that drives expression of a transcription factor (Gal4) in a restricted subset of neurons, including those in R5-R7 (Lacoste et al., 2015; Randlett et al., 2015). When crossed with other transgenic lines that contain an upstream activating sequence (UAS), Gal4 induces selective expression of particular genes useful for visualization, and for chemical or light-mediated manipulation. We first crossed the Tg(-6.7FRhcrtR:gal4VP16) to the Tg(UAS-E1b:Kaede)s1999t line to selectively drive a red fluorescent protein. In addition, we performed these experiments on a transgenic background, Tg(isl1:GFP), that constitutively labeled cranial motoneurons, including extraocular motoneurons, with a green fluorescent protein.

Within R5-R7 (delineated by the rostro-caudal extent of nVII, Figure 1B), we observed expression in ~200 neurons that, in aggregate, comprise a subset of two bilateral vestibular nuclei. The first was the previously characterized utricle-recipient tangential nucleus (Bianco et al., 2012), located adjacent to the ear. The second was the medial vestibular nucleus (Highstein and Holstein, 2006) separated from the tangential nucleus by the lateral longitudinal fasciculus. Figure 1A-1C show the gross morphology of these neurons and their axonal projections to the extraocular motor nuclei. In aggregate, we observed that the axon bundle from these vestibular neurons crosses the midline, ascends rostrally along the medial longitudinal fasciculus (MLF), and projects to extraocular motor nuclei nIII and nIV (Figure 1D-1F).

The utricular vestibulo-ocular reflex utilizes two independent “channels,” or defined neural pathways from peripheral sensation to motor output, to stabilize gaze following pitch and roll tilts. At the level of the extraocular motoneurons, in the larval zebrafish the two channels are segregated along the dorso-ventral axis. First, the ventral-most extraocular motoneurons in nIII project to the inferior oblique (IO) and superior rectus (SR) motoneurons. Together, IO/SR move the eyes up following nose-down pitch tilts. Second, the dorsal-most extraocular motoneurons in nIII project to the inferior rectus (IR), and the dorsally-located nucleus nIV projects exclusively to the superior oblique (SO). Together, IR/SO move the eyes down following nose-up pitch tilts. The somatic organization of nIII and nIV is stable after 5 days post-fertilization (Greaney et al., 2016). Finally, previous electromyographic recordings demonstrates that the SR (nIII) and SO (nIV) muscles are exclusively active during either the nose-down or nose-up phase of pitch-tilts supporting the independence of the two channels (Favilla et al., 1983).

Complementarily, pitch-sensitive vestibular nucleus neurons split into two subtypes, each projecting to only one pair of extraocular motoneurons (Uchino et al., 1982). The first group arborizes exclusively in nIII, innervating IO/SR. The second arborizes in both nIII and nIV, innervating SO/IR. Since nIV is comprised only of extraocular motoneurons that innervate SO, a collateral projection to nIV differentiates vestibular interneurons that respond to nose-up pitch tilts from those that respond to nose-down.

To determine if vestibular neurons labeled in Tg(-6.7FRhcrtR:gal4VP16) comprise both nose-up and nose-down subtypes, we examined their collective projections. We observed that their projection terminated near the ventral-most extraocular motoneurons in nIII (wide view in Figure 1G, close up Figure 1H-1I). The second prominent projection from vestibular neurons goes to extraocular motoneurons in nIV (wide view in Figure 1J, close-up Figure 1K-1L). We conclude that the vestibular neurons labeled in R5-R7 in Tg(- 6.7FRhcrtR:gal4VP16) are poised to respond during both nose-up and nose-down pitch tilts.

To test whether the vestibular neurons labeled in Tg(-6.7FRhcrtR:gal4VP16) projected symmetrically to extraocular motoneurons, we examined the axon collaterals of singly-labeled neurons. To differentiate nose-up from nose-down vestibular neurons, we manually traced the axons of vestibular neurons and used the labeled cranial motor nuclei to categorize their projections, based on the presence/absence of a collateral projection to nIV. We labeled stochastic subsets of vestibular neurons by injecting a plasmid encoding a fluorescent protein into one-cell embryos, Tg(-6.7FRhcrtR:gal4VP16). Experiments were performed on the Tg(isl1:GFP) background to co-label extraocular motoneurons. The majority of labeled neurons (25/27) had only an ascending collateral; the remaining two had a bifurcated axon that both ascended and descended along the MLF. We found that the overwhelming majority (23/27) of labeled vestibular neuron axons had a dorsal collateral projecting to nIV (i.e. nose-up/eyes-down vestibular neurons). One example neuron from the majority population is shown projecting to nIV in Figure 2A-2D and reconstructed as a schematic in Movie M1. In contrast, one example neuron from the minority population, projecting exclusively to nIII with a collateral to the superior rectus motoneurons, is shown in Figure 2E-2G, and reconstructed in Movie M2. Somata of neurons projecting exclusively to nIII were intermingled with those with projections to nIV. By examining labeled neurons at two time points (5 and 11 days post-fertilization) we found that the characteristic collateral projection to nIV in traced vestibular neurons remained stationary (Figure 3).

**Figure 3:**
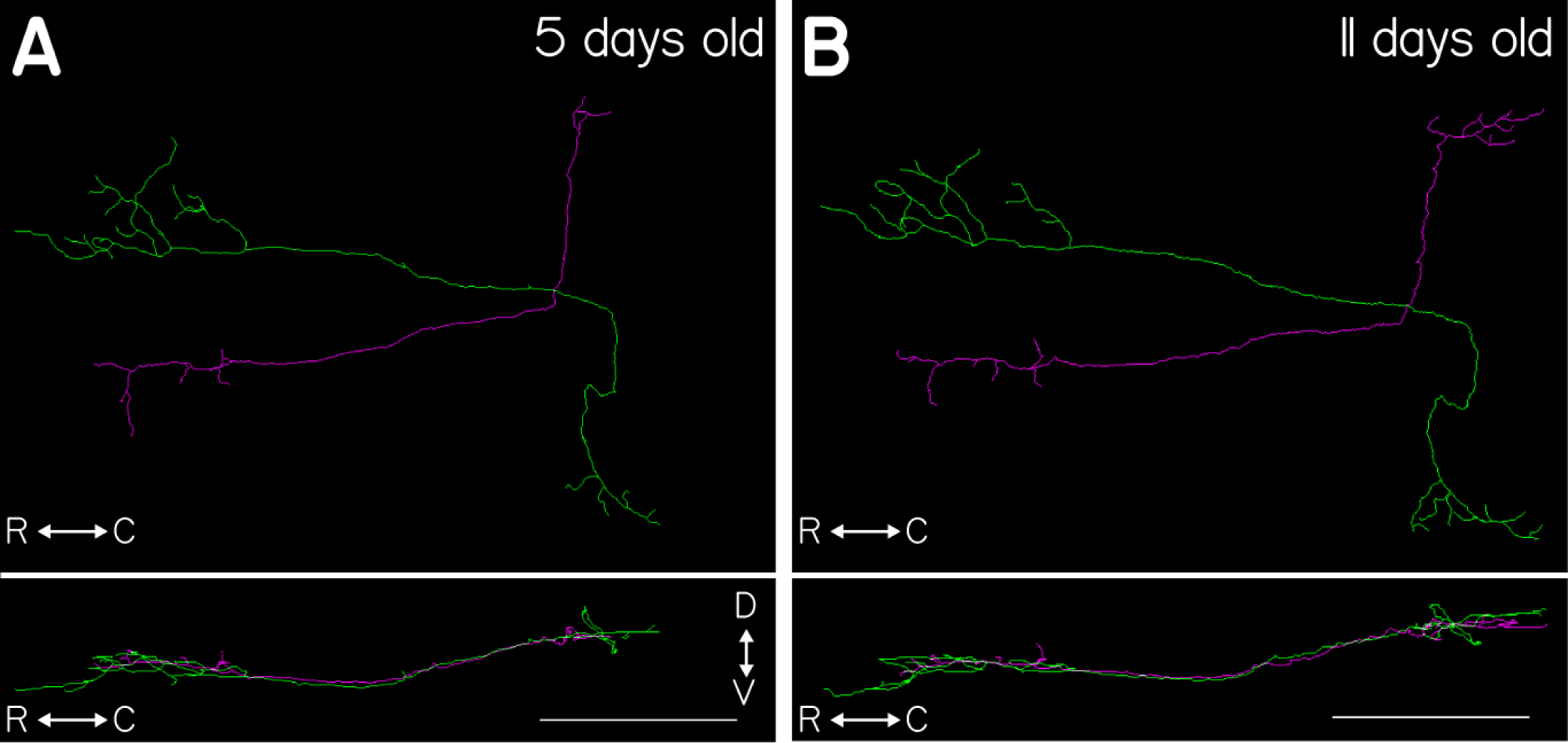
FOR SCREEN: Tracings of two vestibular nucleus neurons from a single fish at two timepoints during development. A, Dorsal (top) and sagittal (bottom) projections of two traced neurons taken from the same fish imaged at 5 days post-fertilization. The magenta trace shows the characteristic projection to the nIV motoneuron pool (magenta arrows) while the green neuron does not. B, Same two neurons traced in the same fish, at 11 days post-fertilization. The same projection to nIV is visible in the magenta tracing (magenta arrow). Scale bars are 100 *μm*.

Our genetically-based labeling technique is limited to neurons within the population labeled in Tg(- 6.7FRhcrtR:gal4VP16). To complement our initial characterization with an unbiased sample of vestibular neurons in R5-R7, we examined the projections of vestibular neurons that had been electroporated with a membrane-targeted fluorescent protein in wild-type animals. Of 20 electroporated animals with singly-labeled neurons in the vestibular nuclei, 15 neurons had an ascending branch along the medial longitudinal fasciculus. 12 of these (80%) had a prominent projection to nIV. Taken together, our data support the conclusion that vestibular neurons in the larval zebrafish project preferentially to extraocular motoneurons that move the eyes down.

To determine whether there was anatomical evidence that the axonal collaterals contained synapses, we labeled presynaptic puncta in Tg(-6.7FRhcrtR:gal4VP16) by crossing to Tg(5xUAS:sypb-GCaMP3) to selectively express a fluorescent protein fused to the presynaptic protein synaptophysin (Nikolaou et al., 2012). We then labeled the extraocular motoneurons by retro-orbital dye fill. We confirmed the presence of presyn-patic puncta proximal to the soma and dendrites of SO and SR motoneurons (Figure 4). Recent expansion microscopy work together with anti-synaptotagmin2b staining confirmed the presence of synaptic puncta between vestibular neurons labeled in Tg(-6.7FRhcrtR:gal4VP16) and extraocular motoneuron somata and dendrites (L. Freifeld and E. Boyden, unpublished observations). These results suggest that the axon collaterals from vestibular neurons labeled in Tg(-6.7FRhcrtR:gal4VP16) to extraocular motoneurons likely contain functional synapses.

**Figure 4:**
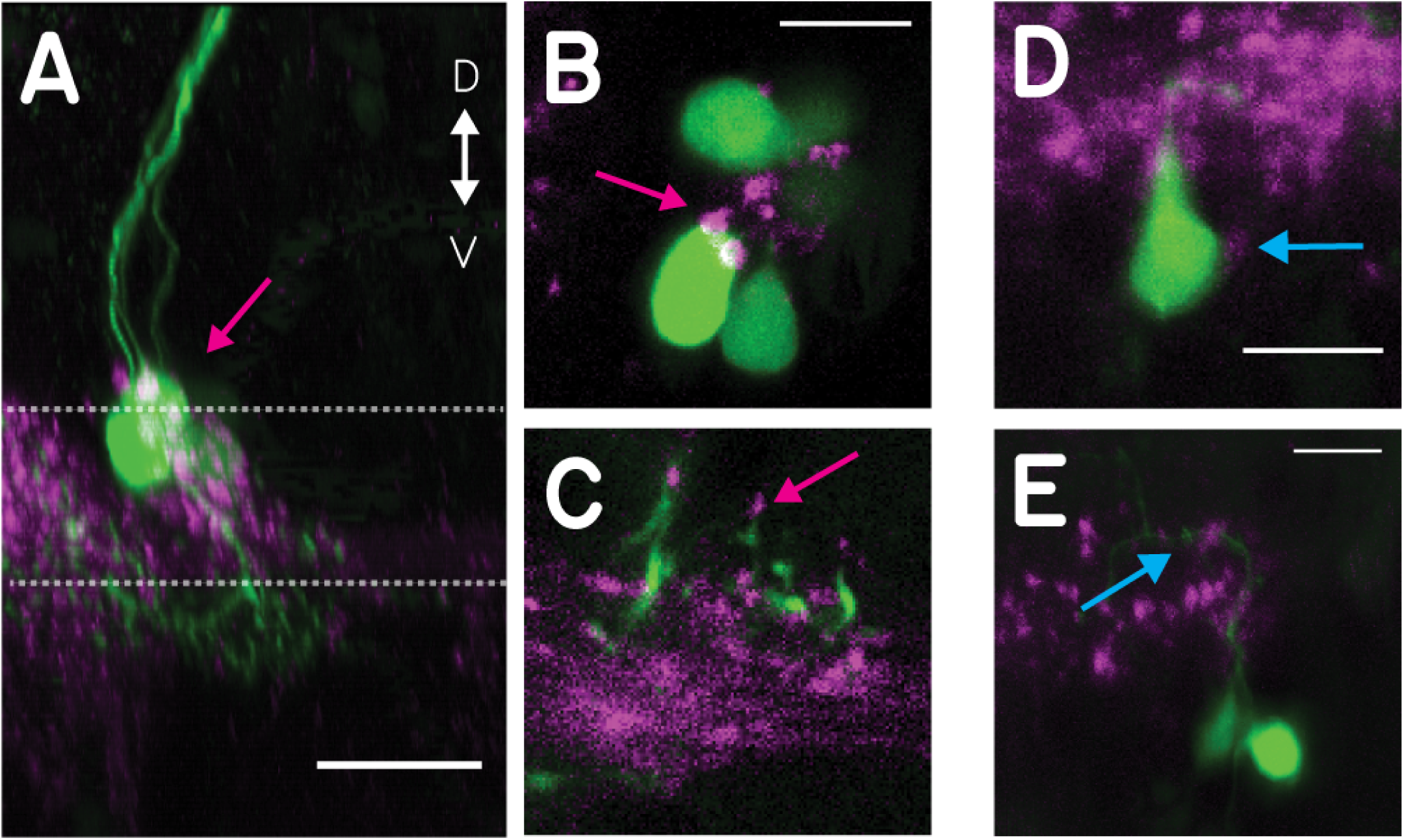
Vestibular nucleus neurons show synaptophysin-positive puncta on their motoneuron targets. A, Sagittal MIP of a labeled SO motoneuron (magenta arrow) in green and the purple synaptic puncta labeled in Tg(- 6.7FRhcrtR:gal4VP16); Tg(5xUAS:sypb-GCaMP3). Dotted lines indicate the planes in 4B-4C, scale 20 *μm*. B,C, Close-up slice of the motoneuron somata in 4A with puncta (magenta arrow), scale 10 *μm*. D, Close-up of a retrogradely labeled SR motoneuron soma (green) with visible purple puncta (cyan arrow), Scale 10 *μm*. E, Close-up of the dendrites of SR motoneurons (green) with visible purple puncta (cyan arrow). Scale 10 *μm*.

### Labeled vestibular neurons are collectively necessary for gaze stabilization following both nose-up and nose-down body rotations

To determine whether the transgenically-labeled vestibular neurons constitute a complete set necessary for both upwards and downwards eye movements following body tilts, we measured gaze stabilization (the vestibuloocular reflex) before and after their removal. We ablated single vestibular neurons individually with a pulsed infrared laser in Tg(-6.7FRhcrtR:gal4VP16). These fish had been crossed to Tg(UAS-E1b:Kaede)s1999t to express a fluorescent protein in vestibular neurons. Further, experiments were performed on the Tg(isl1:GFP) background that labeled adjacent motoneurons in nVII for control ablations (Figure 5A). Following ablation, qualitative observation revealed that horizontal eye saccades and spontaneous jaw movements were present as in normal fish. Ablations eliminated nearly the entire response to body tilts (both nose-up and nose-down): the median decrease in vestibulo-ocular reflex gain was 94.5%±3.5% (n = 14, p = 1.2*10^−4^ Figure 5B). We saw no difference (p = 0.77) in the post-lesion gain for nose-up (0.0165 ±0.0135) and nose-down (0.02±0.0135) body rotations. In contrast, control lesions of somata in the adjacent facial nucleus (nVII) produced no systematic change in the gain (n = 5, 38.5%±24.5%, p = 0.41) or the range (31%±52%, p = 0.44, Figure 5C) of the vestibulo-ocular reflex.

**Figure 5:**
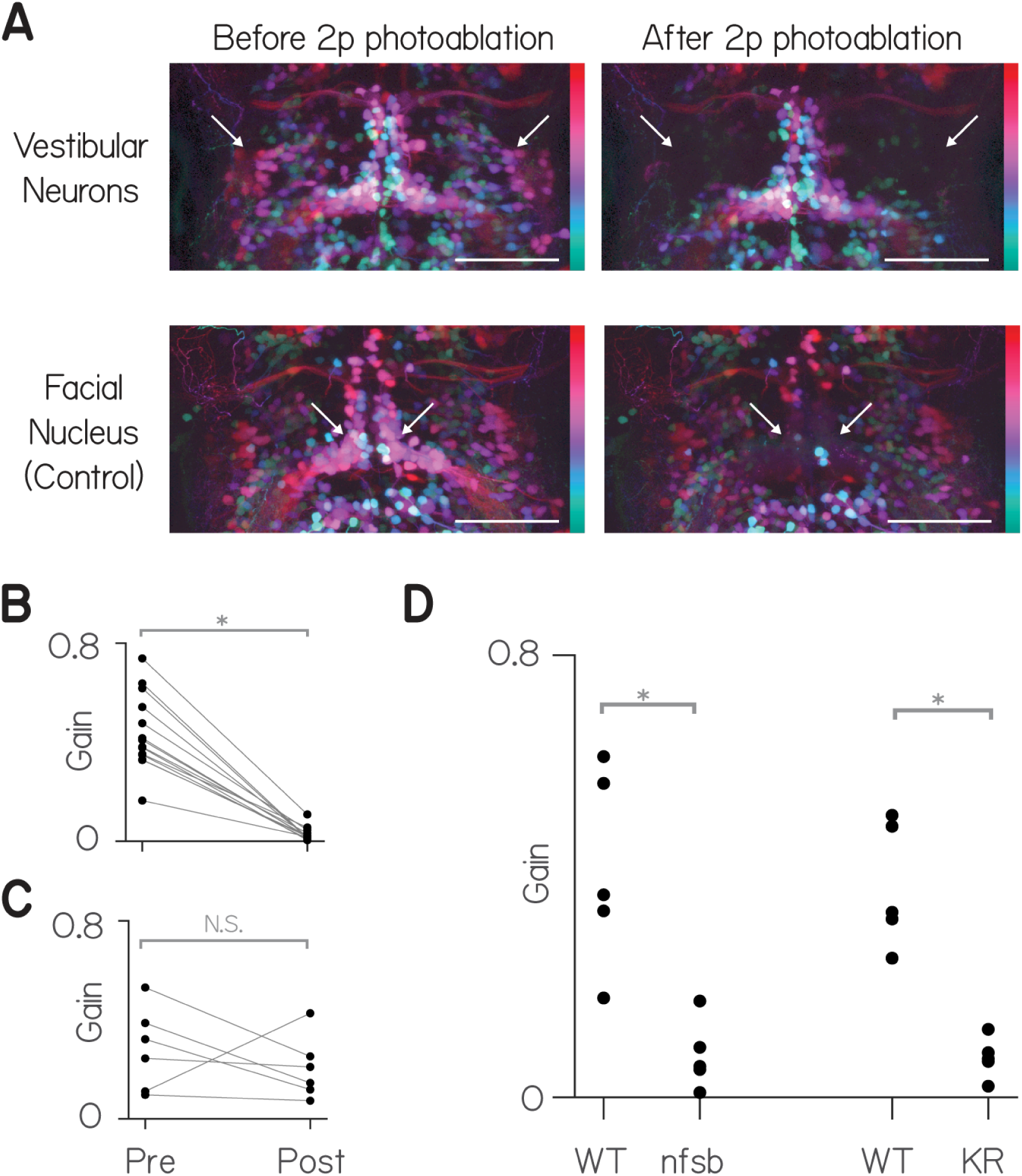
Vestibular nucleus neurons labeled in Tg(-6.7FRhcrtR:gal4VP16) are necessary for both nose-up and nose-down gaze stabilization. A, Transverse MIP of vestibular and control neurons (nVII) in rhombomeres 4-8 in Tg(- 6.7FRhcrtR:gal4VP16); Tg(UAS-E1b:Kaede)s1999t; Tg(isl1:GFP) fish before and after targeted photo-ablation of vestibular neuron cell bodies. Gamma = 0.5 to highlight dim signal, colors indicates depth over ~150*μm*, white arrows point to the general region of targeted cell bodies in either the vestibular nuclei (top row) or the facial nucleus (nVII), scale bar = 150 *μm*. For anatomical localization compare to the right side of Figure 1B. B, Targeted photoablation of vestibular nucleus neurons profoundly impairs the gain of the vestibulo-ocular reflex. C, Control ablations of neurons in the facial nucleus produce no significant change on average to the gain of the vestibulo-ocular reflex. D, Both pharmacogenetic ablation (nitroreductase, “nfsb”) and optogenetic ablation (Killer-Red, “KR”) of vestibular nucleus neurons impair the vestibulo-ocular reflex relative to control siblings.

To confirm the finding that the labeled neurons in Tg(-6.7FRhcrtR:gal4VP16) were necessary for the normal vestibulo-ocular reflex following pitch tilts, we used two additional ablation techniques to target neurons labeled in Tg(-6.7FRhcrtR:gal4VP16). First, by crossing to Tg(UAS-E1b:Eco.NfsB-mCherry) we selectively expressed a protein, nitroreductase (nfsb) that caused neurons to die on exposure to a prodrug, metronidazole (Curado et al., 2007; Pisharath et al., 2007). After exposure to metronidazole, the vestibulo-ocular reflex was significantly impaired in larvae that expressed nfsb compared to their siblings that did not (n=5, p = 0.008, Figure 5C). Next, we crossed Tg(-6.7FRhcrtR:gal4VP16) to Tg(UAS-KillerRed) to selectively express a protein, Killer Red, that causes neurons to die on exposure to green light (Bene et al., 2010). After exposing the hind-brain to green light, the vestibulo-ocular reflex was significantly impaired in larvae that expressed Killer Red compared to similarly exposed siblings (n=5, p = 0.008, Figure 5C). We conclude that vestibular neurons labeled in Tg(-6.7FRhcrtR:gal4VP16) are necessary for compensatory eye movements following either nose-up or nose-down body pitch tilts.

### Labeled vestibular neurons, collectively activated, rotate the eyes down

The circuit that enables correct gaze stabilization following pitch and roll body tilts (Figure 6) permits a specific prediction about the eye movements that might follow collective activation. Three key features of this circuit enable this prediction: 1. two distinct channels selectively sensitive to nose-up and nose-down rotations, 2. excitatory central neurons that all cross the midline, and 3. superior extraocular motoneurons that cross back. Figure 6C-6D show the torsional response to nose-up and nose-down body tilts. There, utricular hair cells in both the left and right ear sense the same pitch tilts. The projection patterns ensure that inputs from a given ear contacts the correct superior eye muscle on the ipsilateral side, and the correct inferior eye muscle on the contralateral side. In contrast, when the fish rolls, both nose-up and nose-down channels ipsilateral to the roll are activated. The two superior muscles are then activated ipsilaterally, while the two inferior muscles are activated contralaterally. In this way, a single circuit can respond appropriately to the two cardinal directions of body rotation sensed by the utricle, the sole source of vestibular sensation in young zebrafish (Beck et al., 2004; Bianco et al., 2012; Mo et al., 2010; Roberts et al., 2017).

**Figure 6:**
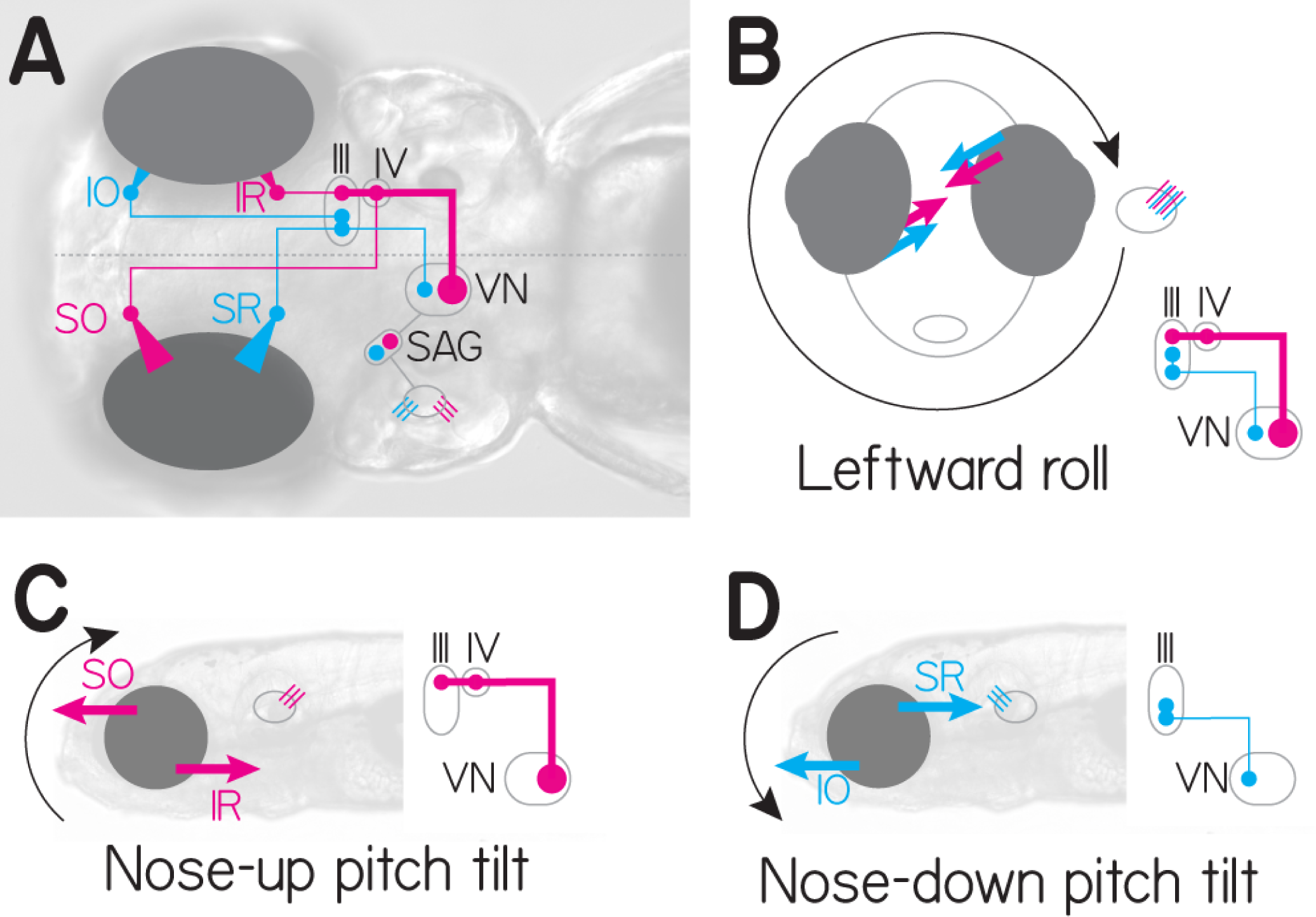
The simplified neural circuit underlying the ocular response to pitch and roll tilts. Cyan = nose down, magenta = nose up channels. A, Wiring diagram of excitatory connections that comprise one hemisphere of the vestibulo-ocular circuit in the larval zebrafish. Utricular hair cells (cyan/magenta) activate neurons in the statoacoustic ganglion (SAG) that project to central vestibular neurons (VN, cyan and magenta). These in turn cross the midline and connect to extraocular motoneuron pools in nIII (SR, IR, IO) and nIV (SO). B, Gaze-stabilization reflex during a roll tilt to the fish’s left. All hair cells in the fish’s left utricle are activated, causing co-contraction of superior (SO/SR) eye muscles ipsilateral to the activated utricle, and inferior (IO/IR) muscles contralateral to the activated utricle. C, Gaze stabilization during nose-down pitch tilts. Select hair cells (magenta) ultimately activate vestibular neurons that project to both nIII and nIV, activating SO (contralateral) and IR (ipsilateral). D, Gaze stabilization during nose-up pitch tilts. Select hair cells (cyan) activate vestibular neurons that project to exclusively to nIII, activating SR (contralateral) and IO (ipsilateral). Note that both left and right utricles are activated by pitch tilts, the ocular response is bilateral.

Collectively activating all utricle-recipient vestibular neurons is therefore equivalent to the fish rolling both left-ward and rightward simultaneously. Consequentially, all four eye muscles on both sides would be expected to contract together. If no eye movement were to result, we would conclude that despite the anatomical asymmetry, the nose-up and nose-down vestibular neuron pools were functionally equivalent. In contrast, a net downward rotation reflects stronger activation of the SO/IR motoneurons (nose-up, Figure 6C) and weaker activation of the SR/IO motoneurons (nose-down, Figure 6D). A net upward rotation reflects the opposite. Any vertical component (SO/SR vs IO/IR) to the eye movement would reflect uneven activation of neurons in the left vs. right hemisphere (Figure 6B), and would be dissociable from the torsional component. We hypothesized that the gaze-stabilization circuit predicts that any systematic eye movement observed along the nose-up/nose-down axis following collective activation of vestibular brainstem neurons must reflect a functional bias in the set of activated neurons.

To determine whether the asymmetry among the population of neurons we observed is functional, we measured eye rotations following collective activation of brainstem neurons labeled in Tg(-6.7FRhcrtR:gal4VP16). We expressed the light-sensitive cation channel, channelrhodopsin-2 (ChR2) and used a fiber-optic cannula (Arrenberg et al., 2009) to target blue light to labeled vestibular neurons in Tg(-6.7FRhcrtR:gal4VP16); Tg(UAS:ChR2(H134R)-EYFP) fish. Since blue light evoked eye movements in wild-type fish (Movie M3), we performed all activation experiments using a blind mutant lacking retinal ganglion cells: atoh7^th241/th241^;

Tg(atoh7:gap43-RFP) (Kay et al., 2001). Strikingly, in every transgenic fish tested, the eyes rotated downward in response to blue light flashes, as if the nose of the fish had moved up. We observed no systematic vertical component to the eye’s rotation. Across fish (n = 10) the amplitude of eye rotation (Figure 7a, black line) scaled with the duration of the light flash, with a peak response of 45°/sec. Crucially, control siblings (n = 3) not expressing ChR2 did not respond to light flashes (Figure 7A, gray line). Laser-mediated ablation of vestibular neurons abolished the light-evoked eye rotation (n=10, Figure 7B). Activation of the population of vestibular neurons is therefore sufficient to rotate the eyes downward, consistent with the asymmetric distribution of anatomical projections.

**Figure 7:**
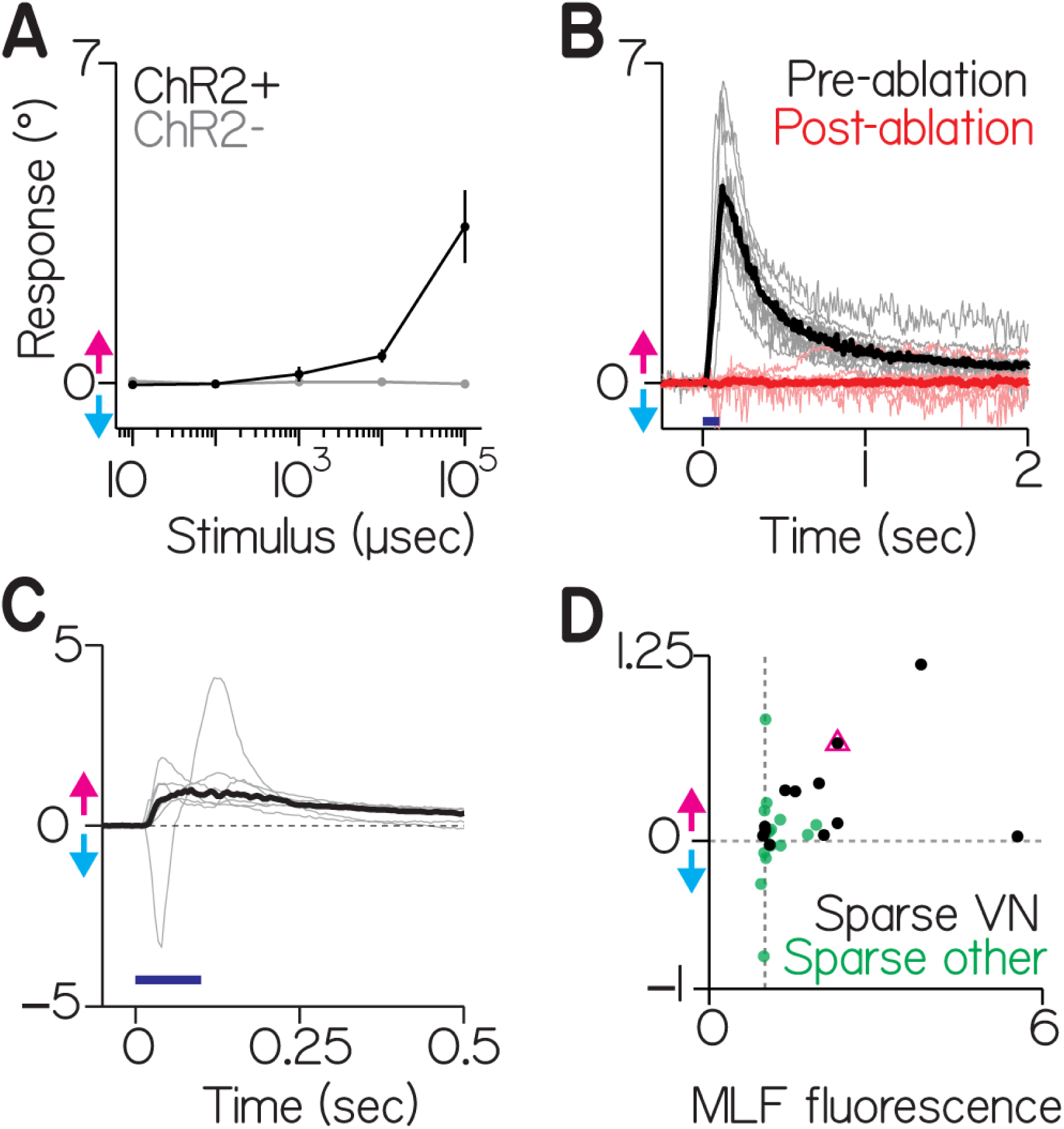
Activating vestibular nucleus neurons generates downward eye rotations. A, ChR2+ fish (black) respond with increasingly large eye movements as the duration of blue light flashes increases. Positive values for eye rotation indicate the direction the eyes would rotate if the nose had been pitched up (magenta arrow), negative values are nose-down (cyan arrow). ChR2siblings do not rotate their eyes in response to blue light (gray line). Points are median ± median absolute deviation. B, Vestibular neurons are necessary for the evoked eye movements. gray lines are individual fish, black lines the median of pre-lesion data, red lines are the same fish post lesion. Blue bar is the duration of the stimulus (100 ms). C, Activation of vestibular neurons in a pan-neuronal fish produces systematic torsional eye rotations in the nose-up direction. Gray lines are the average responses from individual fish with panneuronal expression: the Et(E1b:Gal4-VP16)s1101t; Tg(14xUAS-E1b:hChR2(H134R)-EYFP); atoh7th241/th241 line. Black is the median across fish, blue bar is the duration of the stimulus (100 ms). The single trace with a downward lobe is not due to a torsional movement, but a failure of the eye tracking algorithm to adjust to non-torsional components; video of this fish showing the dominant torsional movement and tracker failure is shown as Movie M4. D, Evoked ocular responses from sparsely labeled fish as a function of the normalized fluorescence in the MLF (a measure of ChR2 expression strength relative to background noise). An MLF fluorescence measure of 1 (gray vertical line) is indistinguishable from background noise, a value of 6 is 6-fold greater than background. Black dots are fish with discriminable vestibular neurons, green dots are fish without (i.e. other labeled neurons). The pink triangle corresponds to the fish whose anatomy is shown in Figure 2A

We extended our test of sufficiency by activating all of the neurons in the region of the vestibular nucleus using a line reported (Scott et al., 2007) to drive expression in all neurons, Et(E1b:Gal4-VP16)s1101t. In all fish tested (n=6), we evoked downward eye rotations in the torsional plane corresponding to nose-up tilts (Figure 7C, Movie M4, note the corruptive horizontal component present in one trace). Both genetically restricted and unbiased activation of vestibular neurons produced net downward eye rotations, and thus the gaze-stabilizing population of vestibular neurons is functionally asymmetric.

To test whether selective activation of vestibular neurons is sufficient to rotate the eyes, and to estimate the variability across neurons, we expressed ChR2 stochastically in subsets of neurons in Tg(-6.7FRhcrtR:gal4VP16) fish on a blind background (atoh7th241/th241; Tg(atoh7:gap43-RFP). Of 27 sparsely labeled fish, 12 had expression in vestibular neurons. As expected from the uneven anatomy, all 12 had neurons with axon collaterals to nIV. Consistent with our categorization of nIV-projecting neurons as “nose-up/eyes-down” we could evoke significant downward eye movements in 10/12 fish (0.23°±0.16°, p < 0.05 relative to baseline for each fish, Figure 7D). Across all fish, the intensity of the projection in the MLF, an estimate of ChR2 expression, predicted the magnitude of the evoked response (Spearman’s rank correlation coefficient = 0.45, p = 0.02, n = 27). These results reveal that subsets of nIV-projecting vestibular neurons are sufficient, but vary in their ability to generate downward eye rotations.

### A simple model shows how biased vestibular populations can better represent nose-up sensations without compromising motor performance

Our data support the hypothesis that labeled premotor vestibular neurons are asymmetrically distributed, over-representing nose-up body tilts, and capable of producing downward eye rotations. To infer whether such an asymmetry might impact motor output and/or sensory encoding, we built a simple model of the synapse between vestibular and extraocular motoneurons. We simulated the ability of differently-sized populations to relay a step in body tilt (encoded by vestibular neuron activity) across a single synapse to produce an eye movement command (encoded by extraocular motoneuron activity). We constrained model parameters and assumptions to reflect known anatomical and electrophysiological properties (Methods). For this model, we assume that the activity of the vestibular neurons is a function of body tilt. We systematically varied two free parameters: the size of the vestibular population, and the number of vestibular neurons that contact a single extraocular motoneuron. As nose-up and nose-down neurons function during distinct phases of pitch tilts (Figure 6) we simulated a single generic population. We evaluated two features of simulated motoneuron activity. First, as a measure of output strength, we report the average activity (reflecting the strength of ocular muscular contraction). Next, as as measure of encoding fidelity, we report the correlation between vestibular input and motoneuron output.

We observed that the magnitude of motoneuron activity could be independent of the number of vestibular neurons upstream (vertical axis in Figure 8C). This dissociation derives from the fact that vestibular neurons encoding nose-up and nose-down body rotations converge on to distinct pools of motoneurons. Consequentially, the key variable that determines the magnitude of motoneuron activity is the number of inputs per motoneuron, not the size of the vestibular population from which it is derived. As expected, increasing the number of vestibular inputs onto a single motoneuron increased its firing rate asymptotically (horizontal axis in Figure 8C). We conclude that when downstream effectors are distinct, as for eye movements, a larger pool of premotor neurons does not necessarily predict differences in the magnitude of motoneuron output. For our system, an asymmetric vestibular circuit could maintain comparable behavioral responses along the eyes-up/eyes-down axis.

**Figure 8:**
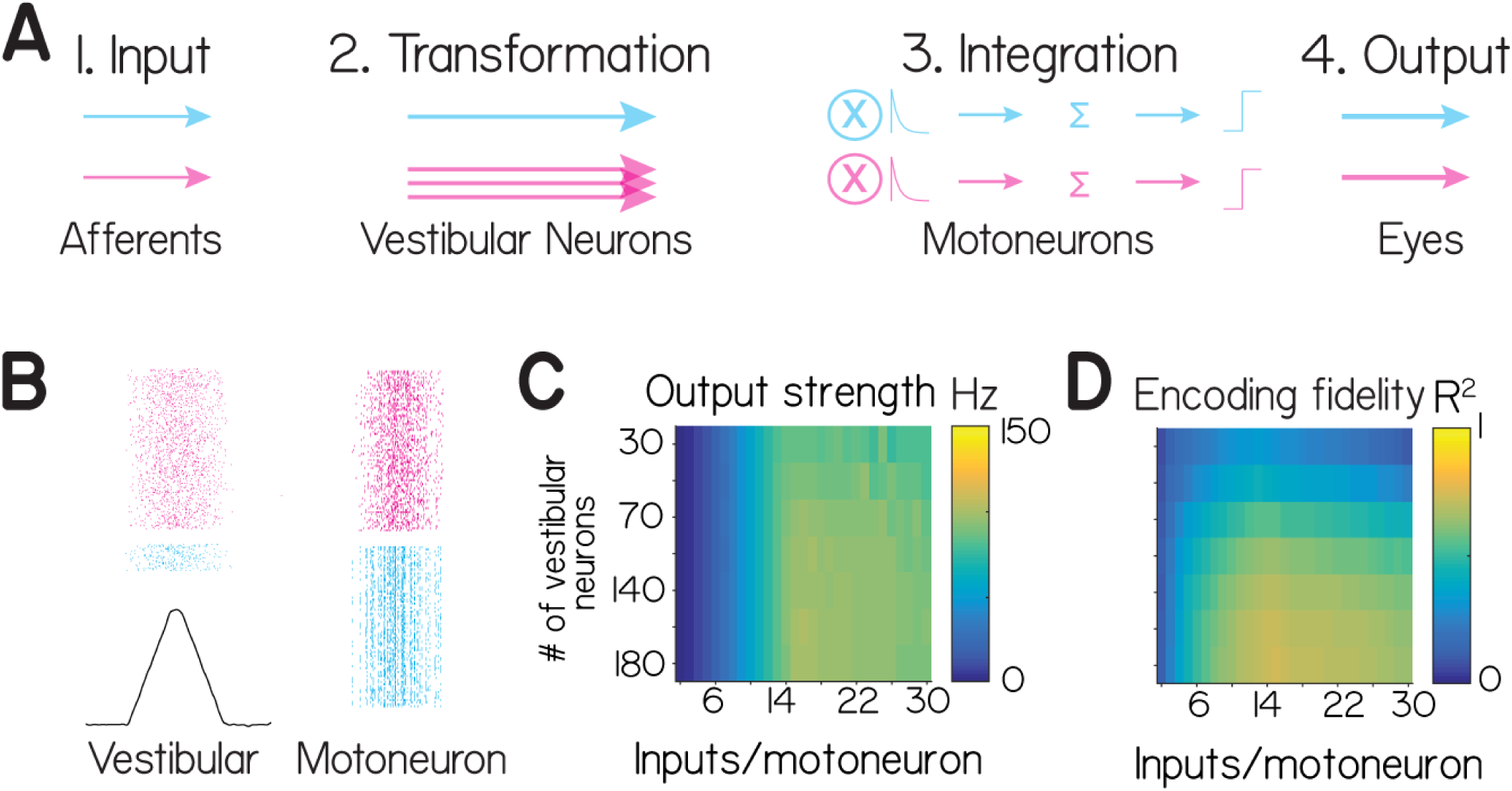
A model of vestibular to extraocular motoneuron transmission. A, Model schematic showing four stages: Input, Encoding, Integration, and Output. They reflect the sensory afferent input, vestibular neuron spiking, synaptic integration and spike generation by ocular motoneurons, and eye movements, respectively. Two channels, nose-up (cyan) and nose-down (magenta) are schematized here as a feed-forward model. We begin with equal inputs to two populations of vestibular neurons, in a 6:1 nose-up to nose-down ratio. Integration consists of convolution, summation, and thresholding. B, One simulation of the model for two different population sizes, 180 neurons (magenta) and 30 neurons (cyan). The first column shows the vestibular neuron activity as a spike raster plot, aligned to the input function in black. The second column shows the motoneuron spikes. For display, half the generated spikes are shown in each raster. C, The “Output strength” (average firing rate) of the post-synaptic neurons as a function of the population size (rows) and number of inputs per motoneuron (columns). D, The “Encoding fidelity” (variance explained, *R*2) in the input rate function by the summed post-synaptic output.

In contrast, we observed that the size of the vestibular neuron pool could impact the ability of motoneurons to represent the dynamics of a step in body position. Temporal structure emerges in the activity patterns of post-synaptic neurons derived from small population sizes (Figure 8b). This similarity across motoneuron activity patterns reflected the coincidence of a limited set of inputs sufficient for a motoneuron spike at a particular time. To test if this limitation constrains the ability of motoneurons to represent the input function, we measured the variance in the input rate function explained by the summed motoneuron activity (*R*^2^). Larger populations were indeed better than smaller populations, and performance varied with the precise number of pre-synaptic inputs (Figure 8D). Adding a basal level of activity equal to 15% of the peak response decreased *R*^2^ but did not change the finding that larger populations were better at representing the input function. We infer from our model that the anatomical asymmetry we observe could permit better encoding of nose-up sensations without compromising gaze-stabilization. If sensory statistics were similarly biased, asymmetric projections from vestibular neurons might therefore be adaptive.

### Premotor vestibular neurons are necessary for a vital and asymmetric postural behavior

To maintain buoyancy, larval zebrafish, whose gills do not yet function (Rombough, 2007), must swim to and maintain a nose-up posture at the water’s surface, where they gulp air, inflating their swim bladder (Goolish and Okutake, 1999). Vestibular sensation is necessary: larval zebrafish without functional utricles fail to inflate their swim bladder and die (Riley and Moorman, 2000). In contrast, vision is not required for this behavior, as blind fish develop normal swim bladders. Gaze-stabilizing vestibular neurons send a second projection to a spinal premotor nucleus, the nucleus of the MLF (nucMLF) (Figure 1), indicating a potential postural role (Bianco et al., 2012).

To test if vestibular neurons are necessary for swim-bladder inflation, we focally ablated vestibular neurons at 72hpf, before fish had inflated their swim bladder, in Tg(-6.7FRhcrtR:gal4VP16);Tg(14xUAS-E1b:hChR2(H134R)-EYFP) fish. We evaluated the fish at 144hpf (Figure 9). Only 1/9 lesioned fish (example in Movie M6) had an inflated swim bladder, compared with 40/42 control siblings (example in Movie M7). To confirm these results, we chemogenetically ablated vestibular neurons at 72 hpf in Tg(-6.7FRhcrtR:gal4VP16);Tg(UAS-E1b:Eco.NfsB-mCherry) fish. As with the targeted lesions, only 1/36 double-transgenic fish inflated their swim bladder and survived, while 36/36 of their non-expressing siblings did. We note that in contrast, fish with post-inflation loss of vestibular neurons (e.g. Figure 5) maintain normal swim bladders. These results define a novel role for vestibular neurons labeled in Tg(-6.7FRhcrtR:gal4VP16) in swim-bladder inflation.

**Figure 9:**
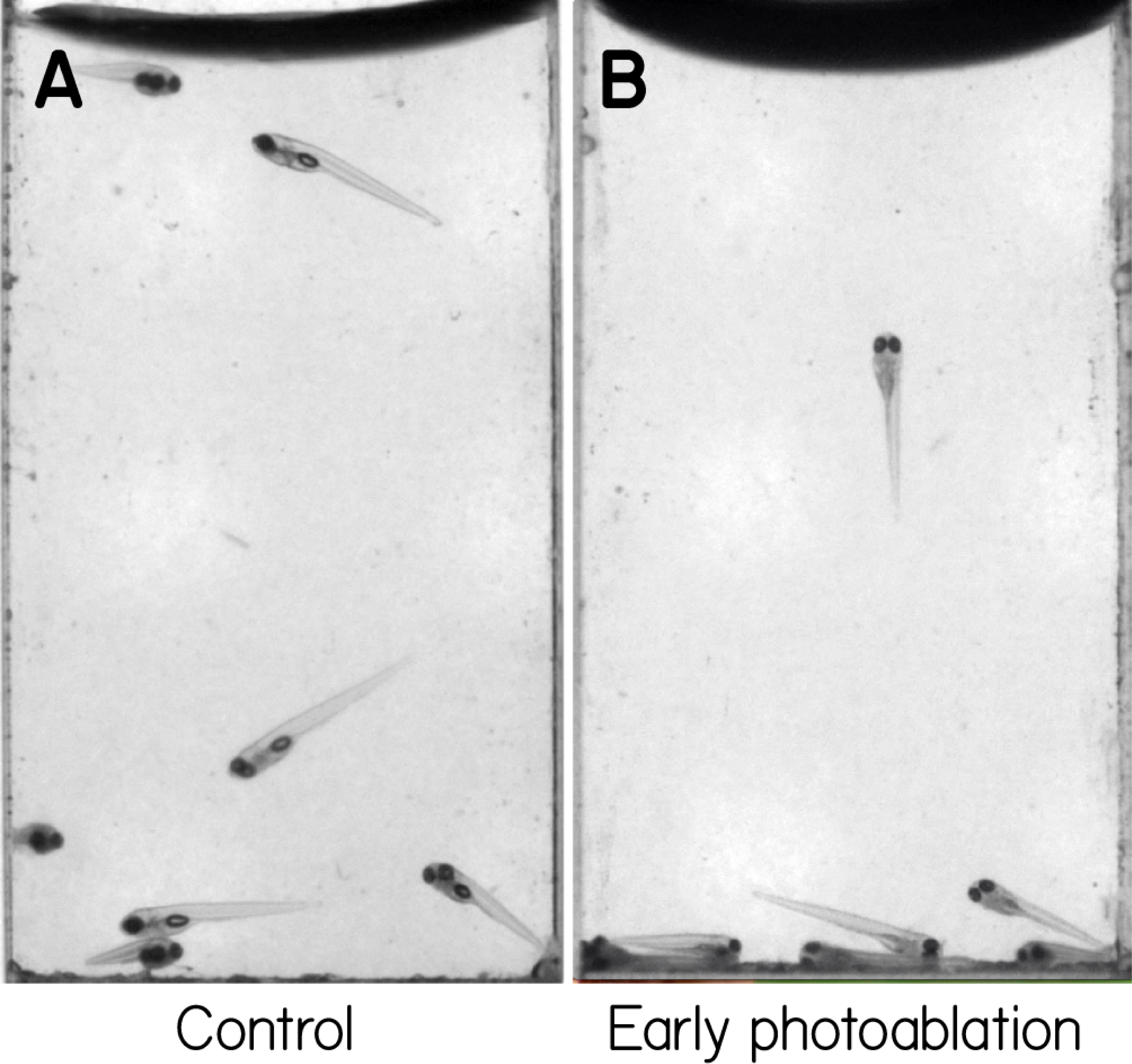
Early ablations of vestibular neurons leave fish unable to inflate their swim bladders. a) Tg(- 6.7FRhcrtR:gal4VP16); Tg(14xUAS-E1b:hChR2(H134R)-EYFP); mitfa -/- fish swimming in a cuvette in the dark at 144hpf. Red arrows point to swim bladders. b) Sibling fish where the vestibular neurons in these fish were photoablated at 72hpf, before swim bladder inflation. Note the absence of a swim bladder, evaluated here at 144hpf. Images are taken from Movies M6-M7.

**Figure 10:**
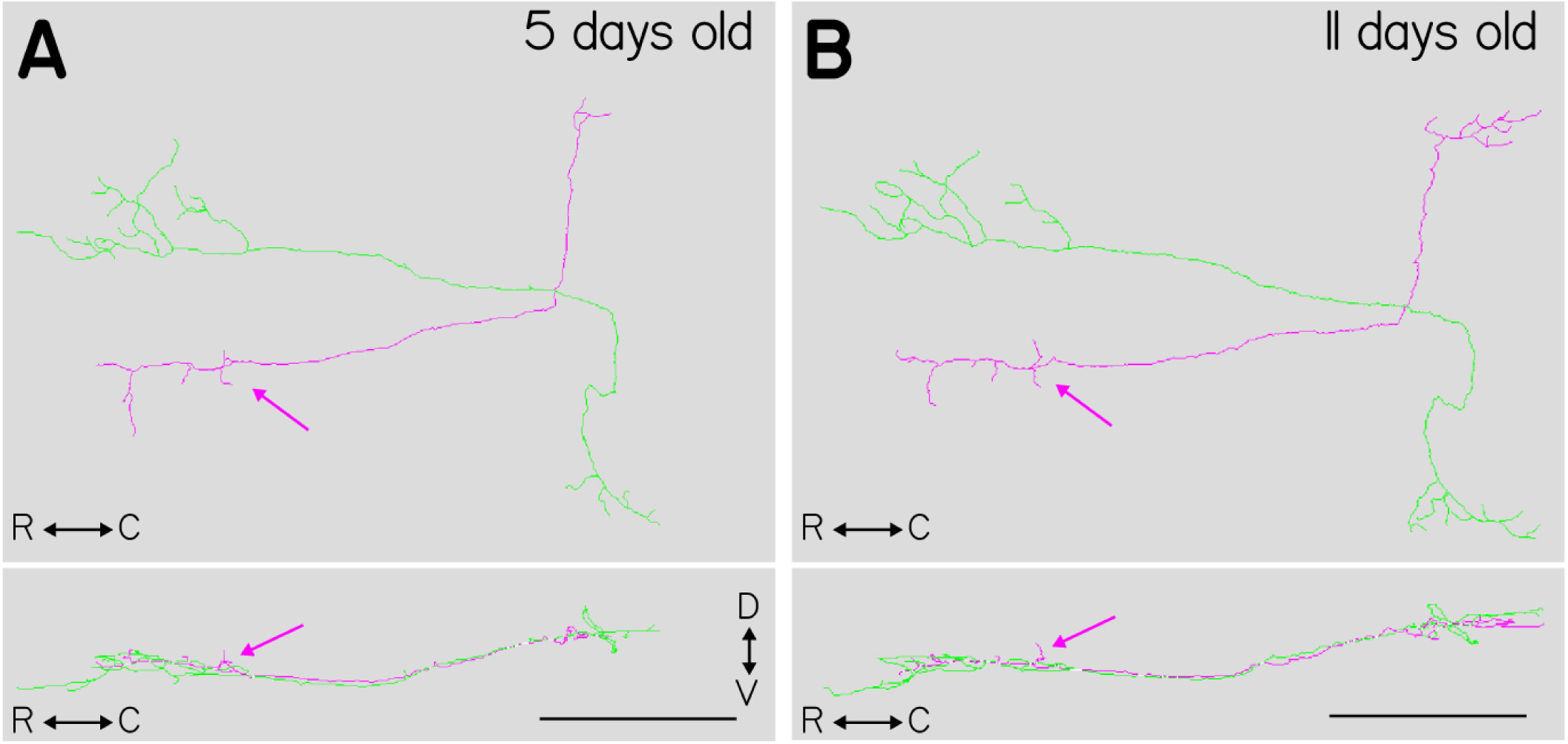
FOR PRINT: Tracings of two vestibular nucleus neurons from a single fish at two timepoints during development. A, Dorsal (top) and sagittal (bottom) projections of two traced neurons taken from the same fish imaged at 5 days post-fertilization. The magenta trace shows the characteristic projection to the nIV motoneuron pool (magenta arrows) while the green neuron does not. B, Same two neurons traced in the same fish, at 11 days post-fertilization. The same projection to nIV is visible in the magenta tracing (magenta arrow). Scale bars are 100 *μm*.

## Discussion

We investigated how the anatomical composition of a genetically-defined population of vestibular interneurons in the larval zebrafish could constrain its function. We first discovered that genetically-labeled neurons project preferentially to motoneurons that move the eyes downward. Ablation of these neurons eliminated the eye movements normally observed following nose-up/nose-down body tilts, establishing their necessity for gaze stabilization. Next, we found that activation produced downward eye rotations, establishing a functional correlate of the anatomical asymmetry. We modeled similar populations with asymmetric projections, and inferred that such architecture could permit better representation of nose-up stimuli while maintaining gaze stabilization performance. Finally, we discovered that early ablation of these neurons impaired swim bladder inflation, a vital postural task requiring nose-up stabilization. Taken together, we propose that preferential allocation of vestibular resources may improve sensory encoding, potentially enabling larval zebrafish to meet ethologically-relevant challenges without compromising behavior.

Our study used a transgenic line, Tg(-6.7FRhcrtR:gal4VP16) to reliably access a genetically defined set of neurons in rhombomeres 5-7 in the medial and tangential vestibular nuclei. The rhombomeric and medio-lateral location of these neurons is consistent with the neurons that receive utricular input in the adult frog (Straka, 2003) and chick (Popratiloff and Peusner, 2007) and comprises a subset of neurons that project to extraocular motoneurons in the larval frog (Straka et al., 2001), juvenile zebrafish/goldfish (Suwa et al., 1996) and chick (Gottesman-Davis and Peusner, 2010). Tg(-6.7FRhcrtR:gal4VP16) does not label neurons within the superior vestibular nucleus in the rostral hindbrain (Cambronero and Puelles, 2000). This absence is notable in light of our ablation experiments that implicate only the neurons labeled in Tg(-6.7FRhcrtR:gal4VP16) as necessary for the torsional vestibulo-ocular reflex. Neurons in the superior vestibular nucleus receive input predominantly from the anterior canal and the lagena (Straka, 2003), and from the anterior canal in monkeys (Yamamoto et al., 1978). Larval zebrafish do not have functional semicircular canals (Beck et al., 2004) nor has the lagena developed (Bever and Fekete, 2002) at the ages we studied here. Therefore, superior vestibular nucleus neurons would not be expected to respond to body rotations, consistent with our observation that the eyes no longer counter-rotate after lesions of Tg(-6.7FRhcrtR:gal4VP16) positive neurons. Further, the superior vestibular nucleus contains predominantly ipsilaterally-projecting, likely inhibitory inputs in adult rays (Puzdrowski and Leonard, 1994), goldfish (Torres et al., 1992, 2008), frog (Montgomery, 1988), rabbit (Wentzel et al., 1995), cat (Carpenter and Cowie, 1985) and monkey (Steiger and Büttner-Ennever, 1979). In the adult goldfish, such inhibitory inputs to extraocular motoneurons were found to be less effective relative to their excitatory counterparts. If the vestibular circuit were similarly constrained in larval zebrafish, it could explain the smaller downward eye movement we saw after collective activation of all neurons. There, the normal downward eye rotation would be compromised, though not eliminated, by inhibition derived from superior vestibular nucleus neurons not labeled in Tg(-6.7FRhcrtR:gal4VP16) but activated in a pan-neuronal line. We therefore propose that the inputs and output of superior vestibular nucleus neurons not labeled in Tg(-6.7FRhcrtR:gal4VP16) render them unlikely to play a major role in the larval zebrafish torsional vestibulo-ocular reflex.

Previous work in larval zebrafish identified the tangential nucleus as the locus of neurons responsible for the utricle-dependent torsional vestibulo-ocular reflex (Bianco et al., 2012). Here, we show similarly profound impairment of the torsional vestibulo-ocular reflex after targeted ablation of a subset of vestibular interneurons in the tangential and medial vestibular nuclei that are labeled in Tg(-6.7FRhcrtR:gal4VP16) larvae. Therefore, we propose that the set of tangential nucleus neurons labeled in labeled in Tg(-6.7FRhcrtR:gal4VP16) are responsible for the utricle-mediated torsional vestibulo-ocular reflex, as those were ablated both here and in (Bianco et al., 2012). Similarly, previous single-cell fills of tangential nucleus neurons revealed three classes of axonal projection neurons: those projecting to the contralateral tangential nucleus, those with a single ascending collateral to nIII/nIV and the nucleus of the MLF and those with both an ascending and descending branch. Ascending and ascending/descending neurons were represented roughly equally (7/16 and 6/16) in the tangential nucleus (Bianco et al., 2012). However, we found that the labeled neurons in Tg(-6.7FRhcrtR:gal4VP16) were almost exclusively of the ascending type (25/27). We therefore propose a further refinement: the neurons responsible for the utricle-mediated torsional vestibulo-ocular reflex are likely the subpopulation of ascending neurons within the tangential nucleus labeled in Tg(-6.7FRhcrtR:gal4VP16). This proposal is consistent with anatomical and functional work in juvenile and adult goldfish, where tangential nucleus neurons with ascending processes were shown to respond to nose-up/nose-down tilts (Suwa et al., 1999) Taken together, our molecular approach permits strong hypotheses that define the essential subset of vestibular neurons responsible for a particular behavior.

The muscles that generate torsional eye movements in fish are responsible for vertical eye movements in front-facing animals (Simpson and Graf, 1981). The behavioral literature is unclear with respect to whether nose-up/nose down gaze-stabilization is asymmetric. Cats may produce stronger downward eye rotations (Darlot et al., 1981; Maruyama et al., 2004; Tomko et al., 1988), but the literature is conflicted as to whether or not such an asymmetry exists in primates: downward (Baloh et al., 1983; Benson and Guedry, 1971; Matsuo and Cohen, 1984) or no biases (Baloh et al., 1986; Demer, 1992; Marti et al., 2006) have both been reported. In foveates, the vestibular brainstem contains the final premotor nuclei for smooth pursuit eye movements. Despite similar abilities to perceive both directions of vertical motion (Churchland et al., 2003), both juvenile and mature monkeys (Akao et al., 2007; Grasse and Lisberger, 1992) and humans (Ke et al., 2013) show a stronger downward response. Our model points a way forward: while the magnitude of the response to comparable stimuli may be comparable, an asymmetric population should better encode dynamic variability, such as experienced in natural settings (Carriot et al., 2014). We propose that characterizing the variation in response to more complex body rotations and target tracking paradigms could uncover behavioral signatures of an anatomically biased circuit.

We found that larval zebrafish do not inflate their swim bladders after early but not late ablation of vestibular neurons. As autonomic neurons are thought to determine swim bladder volume (Smith and Croll, 2011), we propose that the loss of the swim bladder is secondary to postural impairments that follow loss of vestibular neurons labeled in Tg(-6.7FRhcrtR:gal4VP16). In addition to extraocular motor nuclei, these neurons project to the nucleus of the medial longitudinal fasciculus (nMLF) (Bianco et al., 2012). Recent work has established the necessity and sufficiency of spinal-projecting neurons in the larval zebrafish nMLF for postural control and swim initiation (Severi et al., 2014; Thiele et al., 2014; Wang and McLean, 2014). By virtue of their direct projections, and their necessity for swim bladder inflation, we propose that neurons labeled in Tg(- 6.7FRhcrtR:gal4VP16) may affect posture by modulating activity of neurons in the nMLF. As such, our work thus establishes a new molecularly-accessible avenue to explore neural mechanisms underlying postural stabilization.

Zebrafish engage in postural behaviors across their lifespan that are well-suited to nose-up sensory specialization. First, as larvae, they swim along a trajectory dictated by the long axis of their body (Aleyev, 1977). Their bodies are denser than water (Stewart and McHenry, 2010), which ought cause them to sink. Instead, they adopt a nose-up bias to their posture (Ehrlich and Schoppik, 2017), which introduces a vertical component to their swims, enabling them to maintain elevation. Second, larval zebrafish must swim to the surface to gulp air necessary to inflate their swim bladder (Goolish and Okutake, 1999). Finally, most adult teleosts engage in aquatic surface respiration throughout life (Kramer and McClure, 1982), a response to low oxygen saturation that necessitates a continuous nose-up posture at the water’s surface. Our model shows how the anatomical makeup of the vestibular circuits could better encode the nose-up bias in the statistics of behavior. Our findings thus provide a premotor complement to the “efficient coding” framework used to relate the makeup of sensory systems to the statistics of the environment (Simoncelli, 2003).

Asymmetrically organized populations of interneurons are common throughout nervous systems. Asymmetric organization within sensory areas is thought to reflect afferent adaptations (Adrian, 1941; Barlow, 1981; Bendor and Wang, 2006; Catania and Remple, 2002; Hansson and Stensmyr, 2011; Knudsen et al., 1987; Simoncelli, 2003; Xu et al., 2006) but the complexity of most neural circuits makes it challenging to link encoding capacity to adaptive behavior. For asymmetric motor populations, links to behavior are more direct (Esposito et al., 2014; Lemon, 2008; Pasqualetti et al., 2007; Rathelot and Strick, 2009) but the natural sensations that drive these areas are often difficult to define. Our study of vestibular interneurons that play both sensory and premotor roles illustrates how the asymmetry anatomy could better encode nose-up sensations, while maintaining the ability to stabilize gaze. As asymmetric populations of interneurons are common, we propose that other circuits may use similar strategies to meet ethological demands without compromising motor control.

## Author Contributions

Author contributions: DS conceived the study in discussions with IHB, FE and AFS. DS generated the Tg(- 6.7FRhcrtR:gal4VP16) line, designed and built the behavioral apparatus/software, collected, analyzed and modeled data and discussed them with IHB, FE and AFS. IHB performed the focal electroporation and imaging used in Figure 1a. DP and ADD generated the 14xUAS-E1b:hChR2(H134R)-EYFP construct and line. JG and ES helped design, prototype, assemble, troubleshoot and optimize all hardware. DR and JL designed, built, and maintained the 2-photon microscope and control software; DS and DR developed and optimized the lesion protocol. JL and DR made the gal4-VP16 destination vector. DS and AFS wrote the paper.

## Acknowledgements

We thank: Robert Baker for inspiration and extensive insights, Omi Ma for discussions and help with retro-orbital fills, Ian Woods for help with transgenesis, Bill Harris for generously providing the atoh7th241/th241; Tg(atoh7:gap43-RFP) line, Clemens Riegler for maintaining the Tg(5xUAS:sypb-GCaMP3) line, Albert Pan for maintaining the Tg(UAS-E1b:Kaede)s1999t line, the Zebrafish International Resource Center for the Et(E1b:Gal4-VP16)s1101t line, Minoru Koyama for insights into focal lesions, Steve Zimmerman, Karen Hurley, and Jessica Miller for fish care, Bernhard Goetze, Doug Richardson, and Casey Kraft for help with microscopy, Dorothy Barr for library services, Bassem Hassan, Katherine Nagel and the members of the Engert, Schier and Schoppik labs, particularly David Ehrlich, Marie Greaney (who provided the fish schematic in Figure 1), Katherine Harmon, Alix Lacoste, Owen Randlett, and Martin Haesemeyer for helpful discussions. DS was supported by a Helen Hay Whitney Postdoctoral Fellowship and by the National Institute on Deafness and Communication Disorders of the National Institutes of Health under award number K99DC012775 and 5R00DC012775, IHB by a Sir Henry Wellcome Postdoctoral Fellowship from the Wellcome Trust. Research was supported by NIH grants 1R01DA030304 and 1RC2NS069407 awarded to F.E., and R01HL109525 awarded to A.F.S.

## Author Information

Author Information: All raw data, analysis, and figure generation code can be downloaded at http://www.schoppiklab.com/. The authors declare no competing financial interests. Correspondence and requests for reagents or fish lines should be addressed to.

## Movie Legends

### Movie M1: Reconstruction of an nIV-projecting neuron

Data is taken from the same confocal stack shown in Figure 2A-2D. The neuron in shown in black, colored spheres represent the center locus of cell bodies of nIV (magenta) and nIII (cyan) cranial motoneurons. The movie begins with the neuron in a transverse orientation, rotates 90° along the x axis until it is sagittal, and then rotates 90° along the y axis such that the viewer looks caudally down the long axis of the fish towards the tail.

The large projection to nIV is clearly visible. Scale bar is 25 *μm* for all three axes.

### Movie M2: Reconstruction of an nIII-projecting neuron

Data is taken from the same confocal stack shown in Figure 2E-2G. The neuron in shown in black, colored spheres represent the center locus of cell bodies of nIV (magenta) and nIII (cyan) cranial motoneurons. The movie begins with the neuron in a transverse orientation, rotates 90° along the x axis until it is sagittal, and then rotates 90° along the y axis such that the viewer looks caudally down the long axis of the fish towards the tail.

The terminal arbors of this neuron are considerably more restricted than the neuron in M2, and bypass nIV, terminating in nIII. Scale bar is 25 *μm* for all three axes.

### Movie M3: A sample eye movement evoked by a blue light flash in wild-type fish

The left eye of a wild-type fish responding to a flash of blue light. Green box reflects the realtime estimate of the eye’s rotation. Movie is 4 sec, with a 100msec flash after 2 sec. For clarity, the original video was enlarged 4x and slowed down 4-fold (200Hz acquisition, 50Hz playback).

### Movie M4: A sample eye movement evoked by a blue light flash in fish expressing channelrhodopsin in vestibular neurons

The left eye of a Tg(-6.7FRhcrtR:gal4VP16); Tg(14xUAS-E1b:hChR2(H134R)-EYFP); atoh7th241/th241; Tg(atoh7:gap43-RFP) fish responding to a flash of blue light. Gray box reflects the realtime estimate of the eye’s rotation. Movie is 3.6sec long, with a 100msec flash at 1.2sec indicated by a cyan circle.

### Movie M5: A sample eye movement evoked in a fish with pan-neuronal channelrhodopsin

The left eye of a Tg(s1101t:gal4); Tg(14xUAS-E1b:hChR2(H134R)-EYFP); atoh7th241/th241; Tg(atoh7:gap43-RFP) fish responding to a flash of blue light. Green box reflects the realtime estimate of the eye’s rotation.

Movie is 4 sec, with a 100ms flash after 2 sec. For clarity, the original video was enlarged 4x and slowed down 4-fold (200Hz acquisition, 50Hz playback). Note the rapid and transient change in the angle of the green square at the initiation of the evoked eye movement. This reflects the tracker failing due to the nasal-ward component of the eye’s contraction. After the brief failure of the tracker, it recovers, and the downward torsional component of the eye movement becomes visible.

### Movie M6: Fish without vestibular neurons swimming in a cuvette

Tg(-6.7FRhcrtR:gal4VP16); Tg(14xUAS-E1b:hChR2(H134R)-EYFP); mitfa -/- fish swimming in a cuvette. Vestibular neurons in these fish were photoablated at 72hpf, before swim bladder inflation. Note the absence of a swim bladder, evaluated here at 144hpf. These fish are siblings of the fish in Movie M7.

### Movie M7: Fish swimming in a cuvette

Tg(-6.7FRhcrtR:gal4VP16); Tg(14xUAS-E1b:hChR2(H134R)-EYFP); mitfa -/- fish filmed at 144hpf. Note the presence of a swim bladder. These fish are siblings of the fish in Movie M6.

